# Convergent evidence for predispositional effects of brain gray matter volume on alcohol consumption

**DOI:** 10.1101/299149

**Authors:** David AA Baranger, Catherine H. Demers, Nourhan M. Elsayed, Annchen R. Knodt, Spenser R. Radtke, Aline Desmarais, Arpana Agrawal, Andrew C. Heath, Deanna M. Barch, Lindsay M. Squeglia, Douglas E. Williamson, Ahmad R. Hariri, Ryan Bogdan

**Affiliations:** Department of Psychological and Brain Sciences, Washington University, St. Louis, MO; Department of Psychiatry and Behavioral Sciences, Duke University, Durham, NC; Department of Psychology and Neuroscience, Duke University, Durham, NC; Department of Psychiatry, Washington University School of Medicine, St. Louis, MO; Department of Psychiatry and Behavioral Sciences, Medical University of South Carolina, Charleston, SC; Durham VA Medical Center, Durham, NC

## Abstract

**Background:** Alcohol use has been reliably associated with smaller subcortical and cortical regional gray matter volumes (GMVs). Whether these associations reflect shared predisposing risk factors and/or causal consequences of alcohol use remains poorly understood.

**Methods:** Data came from 3 neuroimaging samples (total n=2,423), spanning childhood/adolescence to middle age, with prospective or family-based data. First, we identified replicable GMV correlates of alcohol use. Next, we used family-based and longitudinal data to test whether these associations may plausibly reflect a predispositional liability for alcohol use, and/or a causal consequence of alcohol use. Finally, we evaluated whether GWAS-defined genomic risk for alcohol consumption is enriched for genes preferentially expressed in regions identified in our neuroimaging analyses, using heritability and gene-set enrichment, and transcriptome-wide association study (TWAS) approaches.

**Results:** Smaller right dorsolateral prefrontal cortex (DLPFC; i.e., middle and superior frontal gyri) and insula GMVs were associated with increased alcohol use across samples. Family-based and prospective longitudinal data suggest these associations are genetically conferred and that DLPFC GMV prospectively predicts future use and initiation. Genomic risk for alcohol use was enriched in gene-sets preferentially expressed in the DLPFC and associated with differential expression of *C16orf93*, *CWF19L1*, and *C18orf8* in the DLPFC.

**Conclusions:** These data suggest that smaller DLPFC and insula GMV plausibly represent predispositional risk factors for, as opposed to consequences of, alcohol use. Alcohol use, particularly when heavy, may potentiate these predispositional risk factors. DLPFC and insula GMV represent promising biomarkers for alcohol consumption liability and related psychiatric and behavioral phenotypes.

## INTRODUCTION

Alcohol use and its associated negative consequences are ubiquitous international public health concerns. Worldwide, the average person aged 15 or older consumes 6.2 liters of alcohol annually, and alcohol use accounts for 6% of deaths and 5% of disease burden (1). Combined with the widespread prevalence of problematic alcohol use (e.g., alcohol use disorder lifetime prevalence = 29% (2); current month binge drinking = 26% of US adults (3)), these staggering public health consequences have led to extensive efforts to understand the impact of alcohol use on brain and behavior and identify markers of alcohol use liability.

Neuroimaging studies have consistently shown that alcohol consumption and use disorder are associated with smaller subcortical and cortical gray matter volumes (GMVs), particularly among regions that feature prominently in emotion, memory, reward, cognitive control, and decision making (4–10). While there is evidence that these associations may arise as a consequence of drinking (e.g., reduced neurogenesis in non-human primate models, greater GMV decline among adolescents following initiation of heavy drinking, GMV normalization following abstinence among dependent individuals) (10–17), other data suggest that they may reflect preexisting vulnerabilities that precede and predict drinking initiation and escalating use (18–22).

Here, using neuroimaging data from 3 samples (total n=2,423) (23–25) spanning childhood/adolescence to middle age with prospective or family-based data, we first identify replicable GMV correlates of alcohol use before testing whether these correlates: (1) are plausibly attributable to shared predisposing factors (e.g., shared genetic influence) and/or arise as a consequence from alcohol use, (2) prospectively predict future drinking in young adulthood, and (3) predict drinking initiation in adolescence. Finally, we examined whether genetic risk for alcohol consumption is associated with genes and genetically-conferred differences in gene expression that are preferentially expressed in regions identified by neuroimaging analyses and/or the brain more generally using curated post-mortem data. Here, we applied gene-set enrichment, partitioned heritability, and transcriptome-wide (TWAS)(26) analyses to genome-wide association study (GWAS) summary statistics from the UK Biobank (N=112,117) (27) and AlcGen/CHARGE+ (N = 70,460) (28) studies of alcohol consumption, and RNA-seq data from GTEX (N=81-103)(29) and the Common Mind Consortium (N=452)(30).

## METHODS AND MATERIALS

### Participants

Neuroimaging data were drawn from three independent samples: the Duke Neurogenetics Study (DNS; n=1,303) (23), the Human Connectome Project (HCP; n=897) (24), and the Teen Alcohol Outcomes Study (TAOS; n=223) (25) that assessed a wide variety of behavioral, experiential, and biological phenotypes among young adult college students (DNS), young-middle aged adults (HCP), and children/adolescents (TAOS). The DNS and TAOS samples collected longitudinal data on alcohol use subsequent to the baseline scan. The family-based HCP sample is composed of twin and non-twin siblings. All studies followed protocols approved by relevant Institutional Review Boards and remunerated participants. Additional information regarding each sample is provided in the **Supplement**.

### Alcohol Use Assessment

Alcohol use in the DNS was assessed at baseline (past 12 month use) and follow ups (questions modified to reflect use following the prior assessment) using the Alcohol Use Disorders Identification Test (AUDIT) consumption subscale (AUDIT-C; DNS: α=0.85; M=3.76; SD=2.64; range 0-12) (31, 32). The AUDIT-C was approximated (aAUDIT-C) in the HCP (α=0.786; M=3.42; SD=2.65; range 0-12) and TAOS (α=0.893; M=0.45; SD=1.26; range 0-9) using questions from Semi-Structured Assessment for the Genetics of Alcoholism (SSAGA) (33) and Substance Use Questionnaire (SUQ) (34), respectively. Importantly, the aAUDIT-C for the HCP used maximum lifetime data so that these data could be used to estimate genetic influences. In TAOS, the initiation of alcohol use was defined as attaining a score of 1 or greater on the aAUDIT-C (i.e., participant reports consuming at least one full alcoholic beverage; N=82 initiated during the study; Age: M = 16.68, SD = 1.39, 14.12 – 19.64 yrs). The **Supplement** contains additional details.

### Covariates

The following variables known to be correlated with alcohol consumption/GMV were included as covariates in all analyses: age(35–37), sex(36–39), ethnicity (39, 40), socioeconomic status (SES) (35–39), as well as early-life and recent-life stress(41–43) and intracranial volume (ICV). In samples where multiple scanners were used, scanner was also included as a covariate. In adult samples (DNS, HCP), the presence of any non-substance Axis-I DSM-IV psychiatric disorder was included as a covariate. As the TAOS sample was composed of children/adolescents enriched for a family history of depression, tanner stage and depressive symptoms were included as covariates in all analyses. The **Supplement** contains additional details.

### Magnetic Resonance Imaging Processing

Acquisition parameters and GMV processing for each study are detailed in the **Supplement**.

### Statistical Analyses

#### Discovery

A whole-brain voxel-based morphometry GLM regression analysis was conducted using SPM12 in the DNS sample to test whether alcohol consumption (AUDIT-C) is associated with differences in GMV. Covariates included sex, age, self-reported race/ethnicity (i.e., not-white/white, not-black/black, not Hispanic/Hispanic), scanner id (two identical scanners were used), intracranial volume (ICV), presence of a diagnosis other than alcohol or substance abuse or dependence, perceived stress (PSS), parental education (P-ED), early-life stress (CTQ), and perceived economic status (P-SES). Analyses were thresholded at p<0.05 FWE with a cluster extent threshold of 10 contiguous voxels (k_e_=10) across the entire search volume.

#### Replication

Replication analyses in the HCP sample examined whether lifetime maximum alcohol consumption (aAUDIT-C) predicted differences in GMV only within regions of interest (ROIs) where associations were observed in the discovery DNS sample (Figure 1, **Supplemental Table 1**). ROIs were defined by the AAL atlas (44). A voxelwise GLM regression limited to these ROIs was conducted using multi-level block permutation-based non-parametric testing (FSL PALM v.alpha103; tail approximation p<0.10 with 5,000 permutations), which accounts for the family-structure of the HCP data while correcting for multiple comparisons across space (45–47). Covariates included sex, age, self-reported race/ethnicity (i.e., not-white/white, not-black/black, not Hispanic/Hispanic), intracranial volume (ICV), twin/sibling status (dizygotic/not, monozygotic/not, half-sibling/not), presence of a diagnosis other than alcohol or substance abuse or dependence, perceived stress (PSS), education (ED), and socioeconomic status (SES). Analyses were thresholded at p<0.05 FWE with a cluster extent threshold of 10 contiguous voxels (k_e_=10) across the entire search volume (i.e., across all ROIs collectively).

**Figure 1:**
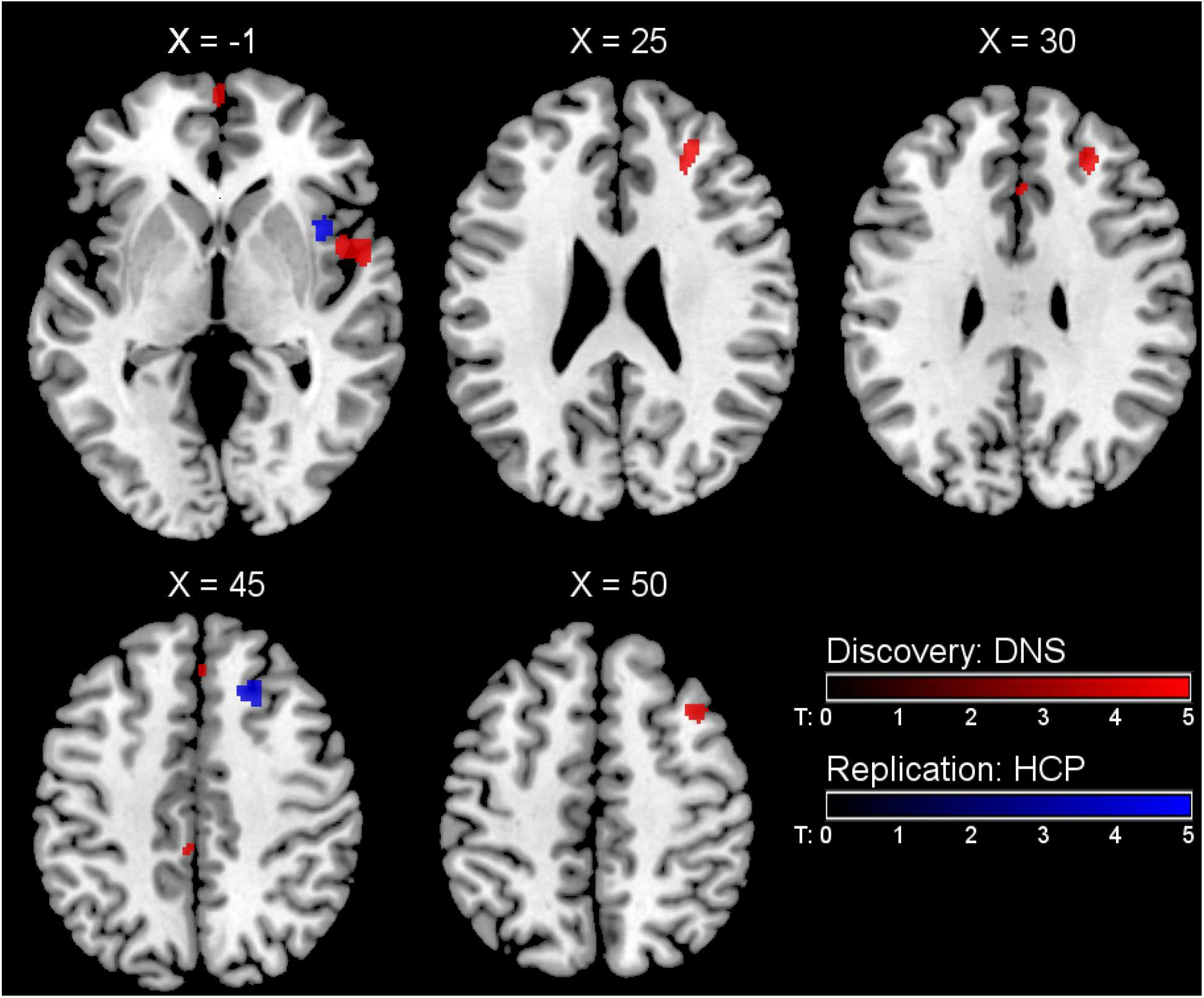
Identification of replicable volumetric associations with alcohol consumption. Statistical parametric map illustrating regions of reduced brain volume associated with increased alcohol consumption (**Supplemental Table 1**), overlaid onto a canonical structural brain image Montreal Neurological Institute coordinates and statistics (DNS: p<0.05, family-wise error whole-brain corrected, ≥10 contiguous voxels; HCP: p<0.05, family-wise error region-of-interest corrected, ≥10 contiguous voxels). Alcohol consumption was not associated with increased volume in any region. Notably, in the HCP dataset, the superior frontal gyrus cluster extended into the right middle frontal gyrus, and was located relatively far (34 mm dorsal) from the original right superior frontal cluster identified in DNS. In contrast, this peak in the HCP was located 11.6 mm away from the right middle frontal peak identified in the DNS. Thus, for the purposes of *post-hoc* analyses, the combined volume of both the right middle and superior frontal gyrus cortices was extracted from both samples. Cluster overlap at an uncorrected threshold and comparison of effect-sizes are shown in **Supplemental Figures 2&3**. DNS = Duke Neurogenetics Study. HCP = Human Connectome Project.

### *Post-hoc* Analyses

Total anatomical ROI GMV of regions associated with alcohol use in both the DNS and HCP (i.e., right Insula and Middle/Superior Frontal Gyrus; see **Results**) were extracted from both datasets for *post-hoc* analyses. Total volumes were used so that effect sizes would not be inflated by selecting only voxels that were specifically associated with the outcome of interest (48).

#### Heritability

SOLAR (Sequential oligogenic linkage analysis routines)-Eclipse software (http://solar-eclipse-genetics.org) (49), in conjunction with the R package ‘Solarius’ (50), was used to assess phenotypic heritability (h^2^; the fraction of phenotypic variance attributable to additive genetic factors), as well as genetic (ρ_g_) and environmental (ρ_e_) correlations (i.e., the fraction of the correlation between two phenotypes that is attributable to either additive genetic or individual-specific environmental factors, respectively) of GMV and alcohol consumption. These analyses were conducted among the subset of related participants from the HCP (n=804; 293 families, 115 MZ and 64 DZ twin pairs and 422 non-twin siblings; excluding singletons and half-siblings). Covariates were identical to neuroimaging analyses. To ensure normality of measurements and accuracy of estimated parameters, an inverse normal transformation was applied to all continuous traits and covariates prior to analyses.

#### Discordant twin analysis

Following evidence that alcohol consumption is co-heritable with volume of the right insula and middle/superior frontal gyri (see **Results**), we examined whether same-sex twin and non-twin sibling pairs discordant for alcohol consumption differed from each other on brain volume in HCP. These analyses examined whether aAUDIT-C was associated with insular or middle/superior frontal volume after accounting for sibling-shared genetic background and experience. Same-sex siblings were considered “high alcohol consumers” or “low alcohol consumers” if their aAUDIT-C score was greater than 0.5 SD above the sample mean (aAUDIT-C > 4.67, or less than 0.5 SD below the sample mean (aAUDIT-C < 1.54), respectively. Concordant sibling-pairs were defined as a pair who were both in the same category of consumption (i.e. high or low), and additionally scored within 1 SD of each other (*low alcohol concordant pairs*: n=117; aAUDIT-C M=0.84, SD=0.77; *high alcohol concordant pairs*: n=54; aAUDIT-C M=0.84, SD=0.77). There were 72 discordant sibling pairs (“low discordant”; aAUDIT-C M=1.25, SD=0.73; “high discordant”; aAUDIT-C M=6.47, SD=1.67). Participants could be included in more than one pair (N=368 individuals) when considering relationships with multiple siblings. Discordancy analyses were conducted using linear mixed models, as sibling pairs are non-independent, using the ‘Psych’ (51) and ‘lme4’ (52) packages in R (53) to account for the multiple-sibling structure within families. Covariates were identical to those used in neuroimaging analyses. Additional information is available in the **Supplement**.

Three contrasts were entered into mixed-effect models, which modeled 3 different possible associations between brain volume, alcohol consumption, and familial/predispositional risk (54). The first tested whether alcohol consumption may plausibly cause reduced brain volume, which would be evidenced by a difference in brain volume between the exposed and unexposed members of discordant pairs. Both the second and third contrasts tested the hypothesis that the association between reduced brain volume and alcohol consumption is driven by a shared predisposition towards both. A non-graded predispositional effect would be evident if discordant pairs – biological siblings who differ in their alcohol consumption – have volumes that do not differ between each other or pairs concordant for high consumption, but are smaller relative to concordant unexposed pairs. On the other hand, evidence of graded liability would be evidenced by equivalent volumes of discordant sibling pairs which are intermediary between concordant high and concordant low users (i.e., significantly higher than concordant high use pairs and significantly lower than concordant low use pairs). **Supplemental Figure 4** provides a graphical depiction of these models.

#### DNS longitudinal changes in alcohol consumption

Hierarchical density-based clustering (R ‘dbscan’ package) (55), was used to detect and remove temporal outlier responses to the follow-up questionnaire (see **Supplement**). The final DNS longitudinal dataset consisted of 1,756 responses from 674 participants, who gave 1-12 (M: 3.59, SD: 2.10) responses, 28-1,034 (M: 350.20, SD: 245.39) days after the baseline visit, between the ages of 18.33 and 23.82 (M: 20.93, SD: 1.24) years (**Supplemental Figure 5**). The R ‘nlme’ package (56) was used to fit a longitudinal multilevel linear model, examining whether GMV predicted AUDIT-C at follow-ups. The ‘nlme’ package was used as it can model different classes of correlation structures between observations. The model included both random intercept and random slope components, with a continuous autoregressive correlation structure. Time was coded as both linear and quadratic age at the date of response (baseline or follow-up). Models tested the interaction between brain volume and age (i.e. does baseline ROI volume predict a different slope of change in drinking behavior as participant’s age?). Covariates were Z-scored, and were identical to neuroimaging analyses, with the addition of second-order interactions between covariates and primary variables (57, 58). Each of the two ROIs were tested in separate models, and p-values were FDR corrected (i.e., 4 tests – middle/superior × linear-age, middle/superior × quadratic-age).

#### TAOS longitudinal initiation of alcohol use

The R ‘lme4 package (59) was used to fit a longitudinal logistic multilevel model, which tested whether baseline brain volume in non-drinking adolescents predicted future initiation of alcohol use. The model included both random intercept and random slope components, and time was coded as both the linear and quadratic age at the date of response. The model tested the interaction between GMV and age (i.e. does baseline ROI volume predict a different likelihood of initiation as participant’s age?). Covariates were Z-scored, and included demographic variables (age, sex, ethnicity, and SES), stress (CTQ and SLES), tanner-stage, MFQ-scores, family history of depression, age at MRI scan, and intracranial volume. Second-order interactions between covariates and primary variables (e.g., Middle Frontal volume × Sex, Middle Superior volume × SES, Age × Sex, Age × SES, etc.) were also included (57). Each of two ROIs were tested in separate models - right superior frontal cortex and right middle frontal cortex. Insula volume was excluded as it was not significant in DNS longitudinal analyses. P-values were subsequently FDR corrected (4 tests).

#### SNP-Based Enrichment

We tested whether the SNP-based heritability of alcohol consumption is enriched in brain-expressed gene-sets, and whether this enrichment is specific to any region. Stratified LD-score regression (60–62) was applied to summary statistics from the genome-wide association study of alcohol consumption in the UK Biobank (N=112,117) (27). Tissue-enriched gene-sets, provided by the Alkes Group, were generated using data from the GTEx Consortium (29). In this analysis, a gene is assigned to a gene-set if it shows greater enrichment in that tissue than 90% of genes. Gene-sets for brain regions were generated both by comparing each region to all non-brain tissues, and by comparing each brain region to all other regions. It was further tested whether genetic associations with alcohol consumption are enriched in brain-expressed gene-sets using MAGMA (63), implemented through FUMA (64).

#### Transcriptome-Wide Analysis (TWAS)

Following evidence that genetic associations with alcohol consumption are enriched in brain-expressed gene-sets, and that the SNP-based heritability of alcohol consumption is enriched in these gene-sets, we tested whether genetic risk for alcohol consumption is predictive of changes in post-mortem gene expression in regions identified in our neuroimaging analyses and the human brain more generally. Pre-computed gene-expression RNA-seq weights for nine brain regions and the liver from GTEX (29) were analyzed using the FUSION suite (26). We tested whether genetic risk for alcohol consumption, as determined by the results from the UK Biobank GWAS (27), is associated with differential RNA expression. Results were bonferroni-corrected for n=9,839 tests across the ten tissues. Replication was tested using independent GWAS data from an earlier non-overlapping study of alcohol consumption (N = 70,460) (28) and computed gene-expression weights for the dorsolateral prefrontal cortex from the CommonMind Consortium (30). As the gene that showed the strongest association in the discovery dataset was not present in the replication data, we examined whether any of the gene-expression associations at p-fdr<0.05 in Brodmann Area 9 were significant in the replication data (see **Results**). Any identified genes, were probed for association with other phenotypes in prior GWAS using PheWAS within the GWAS ATLAS (http://atlas.ctglab.nl/). Brodmann area 9 in the GTEx dataset and DLPFC in the CommonMind consortium dataset overlap with the prefrontal regions implicated in our neuroimaging analyses (i.e., superior and middle frontal gyri; see **Results**). No postmortem insula data were available.

## RESULTS

Whole brain discovery analyses in DNS revealed that greater alcohol consumption is associated with lower GMV across 8 clusters (Figure 1; Table 1), encompassing regions identified in prior studies of unselected samples (5, 17) and among individuals with alcohol use disorder (4, 6). The associations with two of these clusters (right insula, right superior/middle frontal gyrus) replicated within an ROI analysis in the HCP (Figure 1; Table 1). Statistics are presented in **Tables** and **Figure** legends^1^.

**Table 1.**
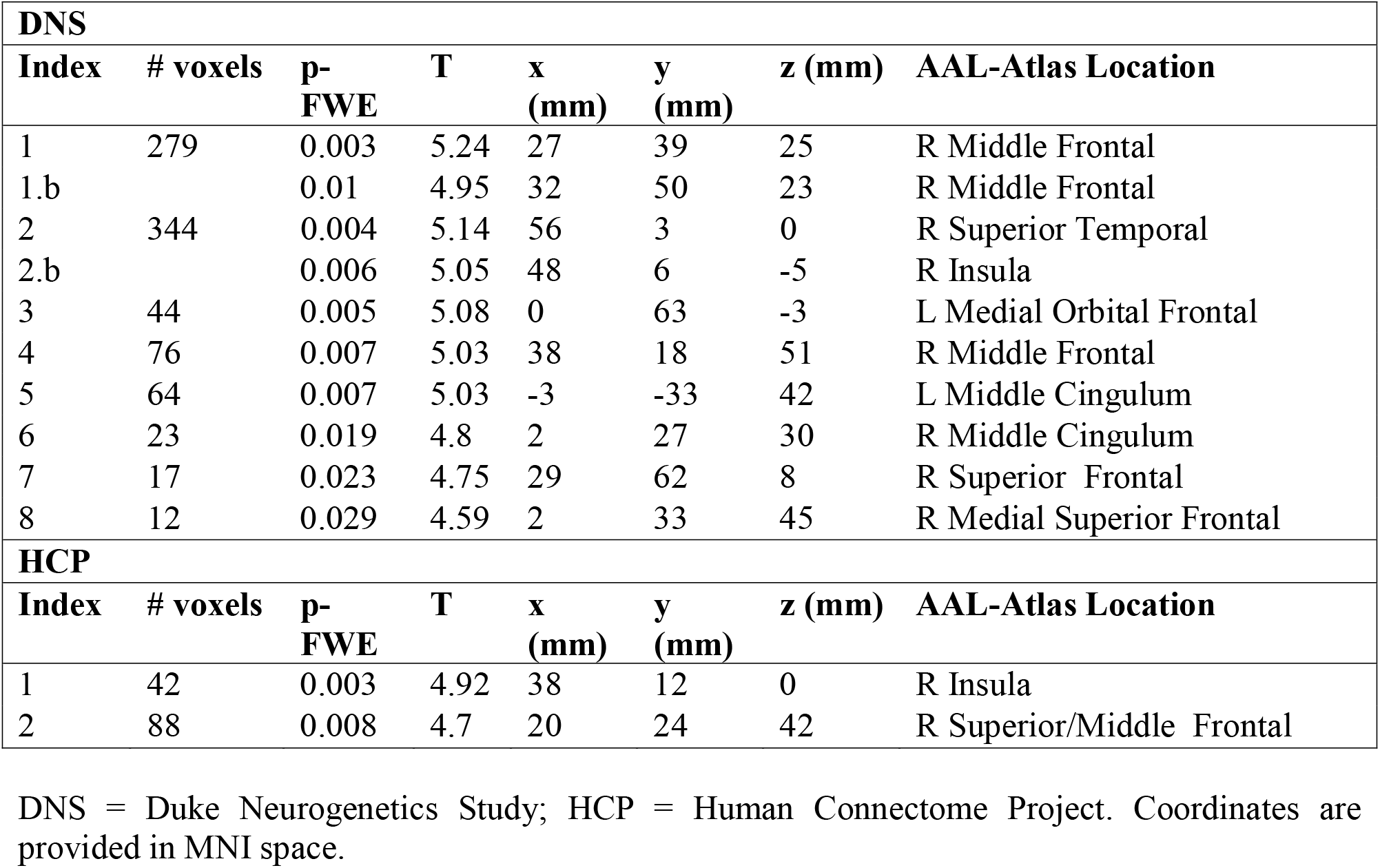
Location of volumetric reductions associated with alcohol consumption

Family-based analyses in the HCP (N=804) revealed that alcohol consumption and GMV of the right insula and right middle/superior frontal gyrus are moderately to largely heritable (Figure 2A; **Supplementary Table 1**). Moreover, decomposition analyses showed that phenotypic correlations between frontal and insular GMV and alcohol consumption are attributable to shared genetic, but not environmental, influences (Figure 2B; **Supplementary Table 1**). Analyses within twin and sibling pairs in the HCP concordant or discordant for the extent of lifetime alcohol use revealed that, relative to siblings concordant for low alcohol use, siblings concordant for high use or discordant for use (i.e., one high use, one low use) had lower insular and frontal gray matter volumes (Figure 2C&D; **Supplementary Table 2**). Further, GMVs did not differ between low and high alcohol-using members of discordant pairs. As shared genetic and familial factors are matched within pairs, this pattern of results suggests that smaller frontal gyri and insula GMVs may reflect preexisting vulnerability factors associated with alcohol use, as opposed to a consequence of alcohol use.

**Figure 2:**
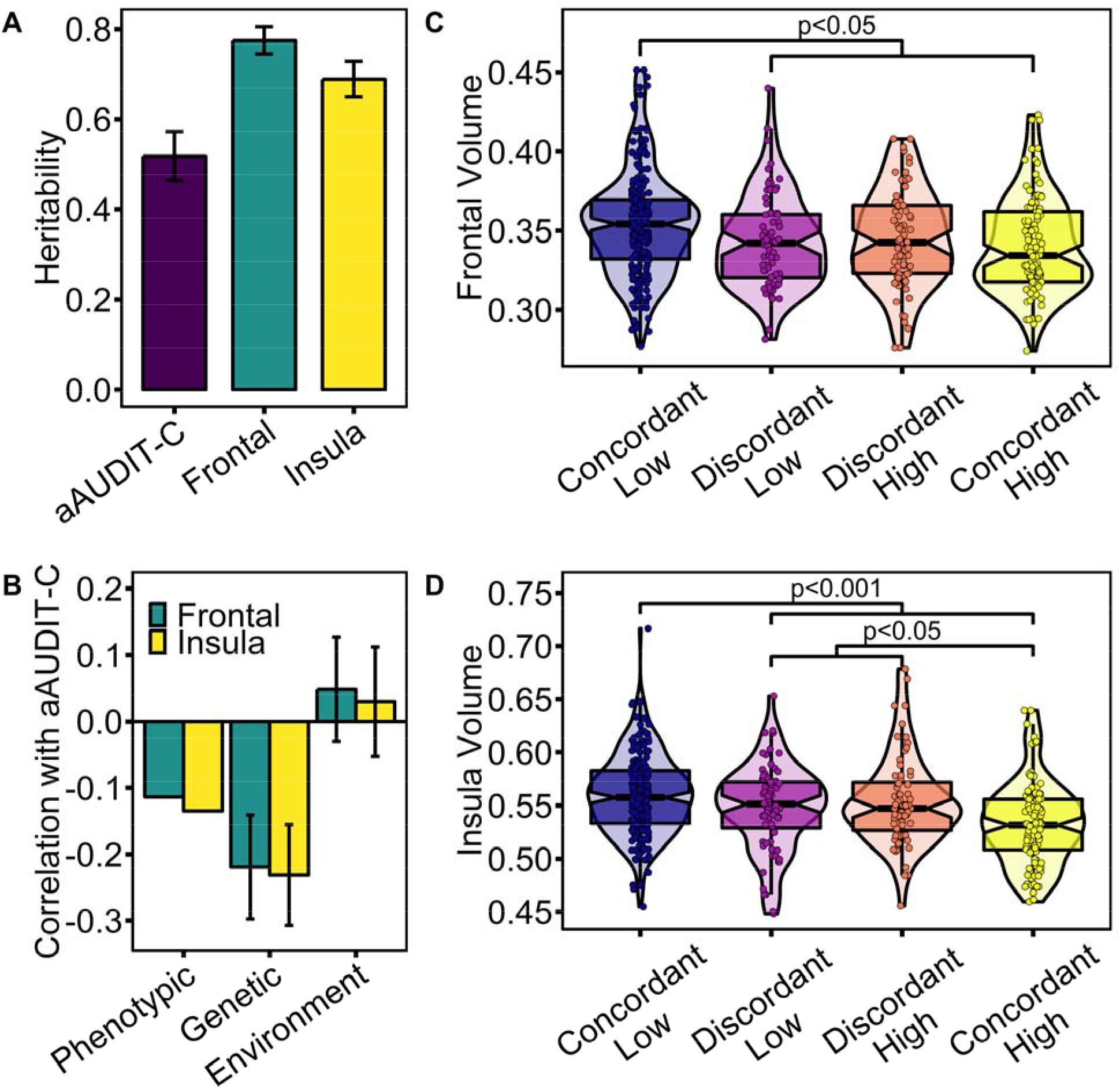
Shared genetic predisposition between alcohol consumption and brain volume. **HCP: A)** Alcohol consumption scores (aAUDIT-C) and gray-matter volume of the right insula and right middle/superior frontal cortex were all observed to be heritable (aAUDIT-C: 51.79%, p<2.2×10^−16^; insula: 68.83%, p<2.2×10^−16^; frontal: 74.46%, p<2.2×10^−16^; **Supplemental Table 1**). **B)** Significant phenotypic correlations between aAUDIT-C scores and volumes of the right insula and middle/superior frontal gyri are attributable to shared genetic (Insula: −0.2314, p=0.0022; Frontal: −0.2192, p=0.0054), but not environmental factors (**Supplemental Table 1**). **C&D)** Distribution of (**C**) right insula and (**D**) right middle/superior frontal volumes by alcohol exposure group. High = aAUDIT-C score > sample mean + 0.5 SD (i.e. > 4.67); Low = aAUDIT-C score < sample mean - 0.5 SD (i.e. < 1.54); Concordant = both siblings are in same group; Discordant = one sibling is High, while other is Low. Contrast comparisons found evidence for predispositonal effects of brain volume on alcohol consumption in both cases (Insula: Graded Liability: β=-0.0037, p=0.049, Predispositonal: β=0.0037, p=0.0006; Frontal: Predispositonal: β=0.0019, p=0.029; **Supplemental Table 2**).

Using available longitudinal data from the DNS (N=674), we found that lower GMV of the right frontal gyri, but not insula, predicted increased future alcohol consumption, over and above baseline consumption, but only in individuals who are under the legal age of drinking (i.e., younger than 21) in the United States (Figure 3A; **Supplementary Table 3**). Similarly, in the TAOS longitudinal sample of children and adolescents, lower right middle and superior frontal gyrus GMV predicted the initiation of alcohol use at an earlier age in those who were nondrinkers at baseline (Figure 3B&C; **Supplementary Table 4**).

**Figure 3:**
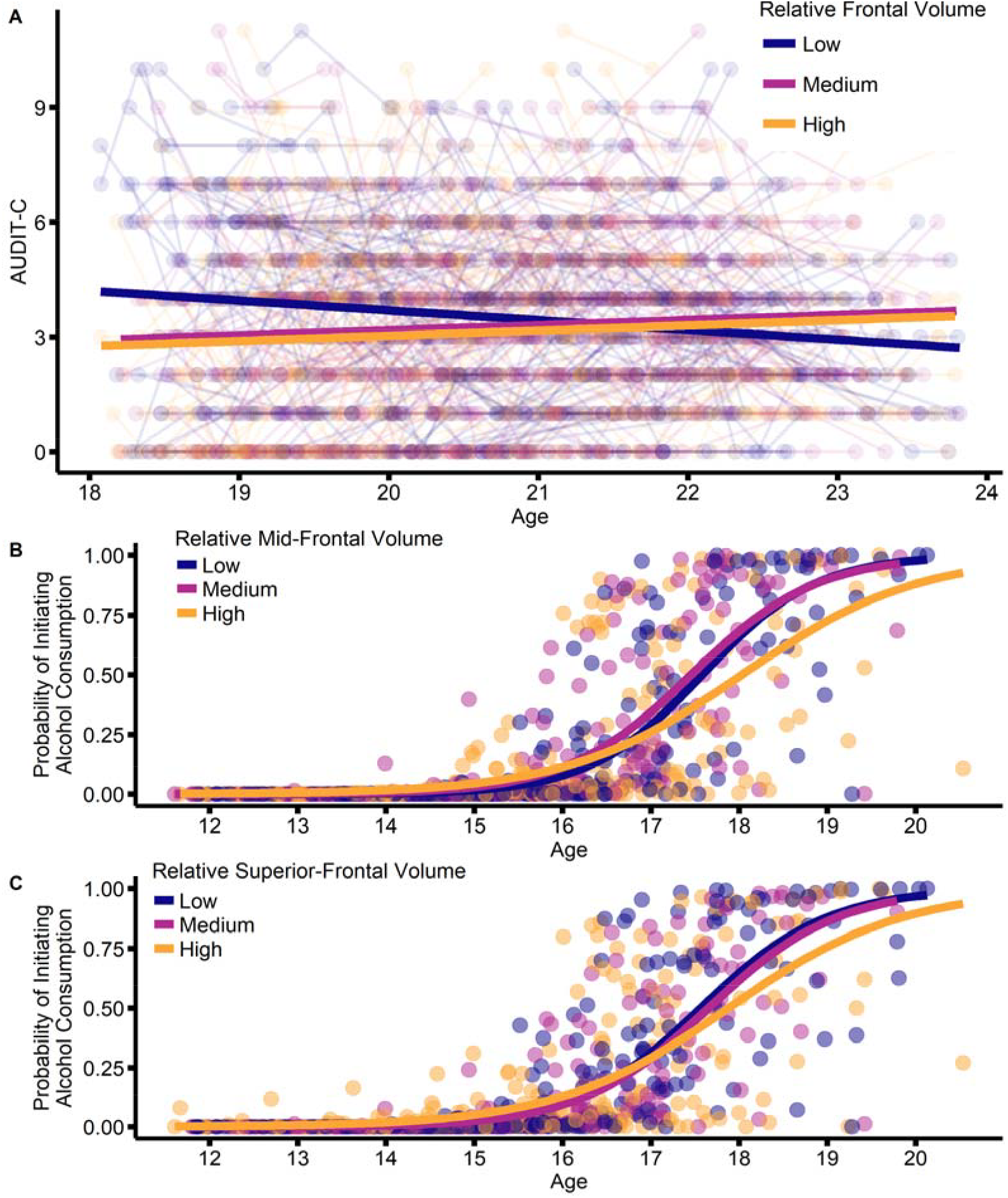
Frontal volume prospectively predicts alcohol use and initiation of consumption. **A)** DNS: Participants with reduced volume of the right middle/superior frontal cortex reported elevated alcohol consumption before the age of 20.85 years following the neuroimaging scan, and after accounting for baseline drinking (Frontal × Age interaction: β=0.150, p-fdr=0.008; **Supplemental Table 3**). **B&C**) TAOS: Participants with increased volume of the right middle and superior frontal cortex report initiation of alcohol consumption at an older age (Mid-Frontal × Age interaction: β=-57.042, p-fdr=0.036; Superior-Frontal × Age interaction: β=-60.74, p-fdr=0.036 **Supplemental Table 4**). Analyses were conducted with continuous data; partitioned into three equally-sized groups according to volume was done for display-purposes only.

Gene-based association and partitioned heritability enrichment analyses of the UK Biobank GWAS of alcohol consumption revealed enrichment only among brain gene-sets. Moreover, Brodmann Area 9, which overlaps with the frontal region in which we observed a replicated negative association between GMV and alcohol consumption that is attributable to shared genetic influence and predictive of drinking initiation, was among the regions with strongest enrichment (Figure 4, **Supplemental Data**). A transcriptome-wide association study (TWAS) analysis of these GWAS data similarly found that genetic risk for alcohol consumption was significantly associated with differences in gene expression across the brain within the GTEx dataset, including *C16orf93* within Brodmann Area 9 (Figure 5; Table 2). Other identified genes include *CWF19L1* in the putamen and cortex, *AC074117* in the cerebellum, and *C1QTNF4* in cortex (Figure 5). *C16orf93* was not available in the TWAS replication dataset (i.e., (28) and the CommonMind Consortium (30) dataset; Table 2). Three additional genes (i.e., *CW19L1*, *PHBP9*, *C18orf8*) were associated with genetically-related differential expression in Brodmann Area 9 that survived FDR correction in the discovery dataset (i.e., across all tissues tested), two of which (i.e., *CW19L1* and C18orf8) were available in the TWAS replication dataset (Figure 5; Table 2)^2^. Genetically-related expression of *CWF19L1* and *C18orf8*, within the DLPFC of our TWAS replication dataset were related to genomic risk for alcohol consumption in the same direction as observed in the discovery dataset, and survived fdr-correction for multiple comparisons (Table 2). Notably, genetic risk for alcohol consumption was not significantly associated with the expression of any gene in the liver (Figure 5). PheWAS using the GWAS ATLAS revealed evidence that both *CWF19L1* and *C18orf8* have been implicated in a host of phenotypes, including psychiatric conditions and related traits such as executive function, schizophrenia (*CWF19L1*), and substance use (*C18orf8*; **Supplemental Data**).

**Figure 4:**
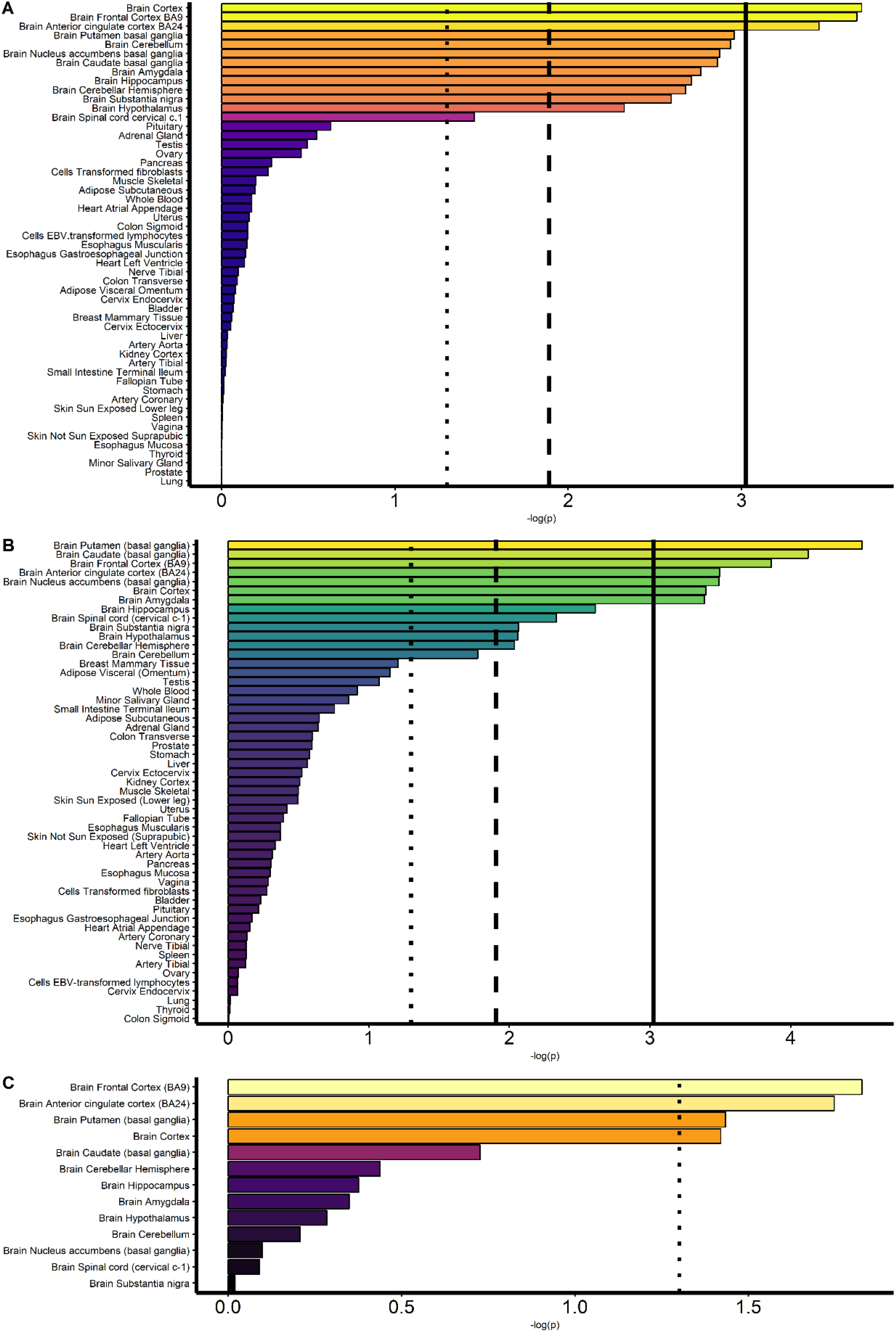
Tissue-specific Enrichment of Alcohol Consumption GenomicRisk. Enrichment of alcohol consumption GWAS (UK Biobank, N=112,117) **A)** associations and **B&C)** heritability, in gene-sets defined by the relative expression of genes across **A&B)** all tissues, and **C)** within the brain, in the GTEX data set (**Supplemental Data**). X-axis and color-scale represent the significance of the enrichment (-log-scale of the p-value). Solid, dashed, and dotted lines represent Bonferroni-corrected, FDR-corrected, and nominally significant p-values, respectively.

**Figure 5:**
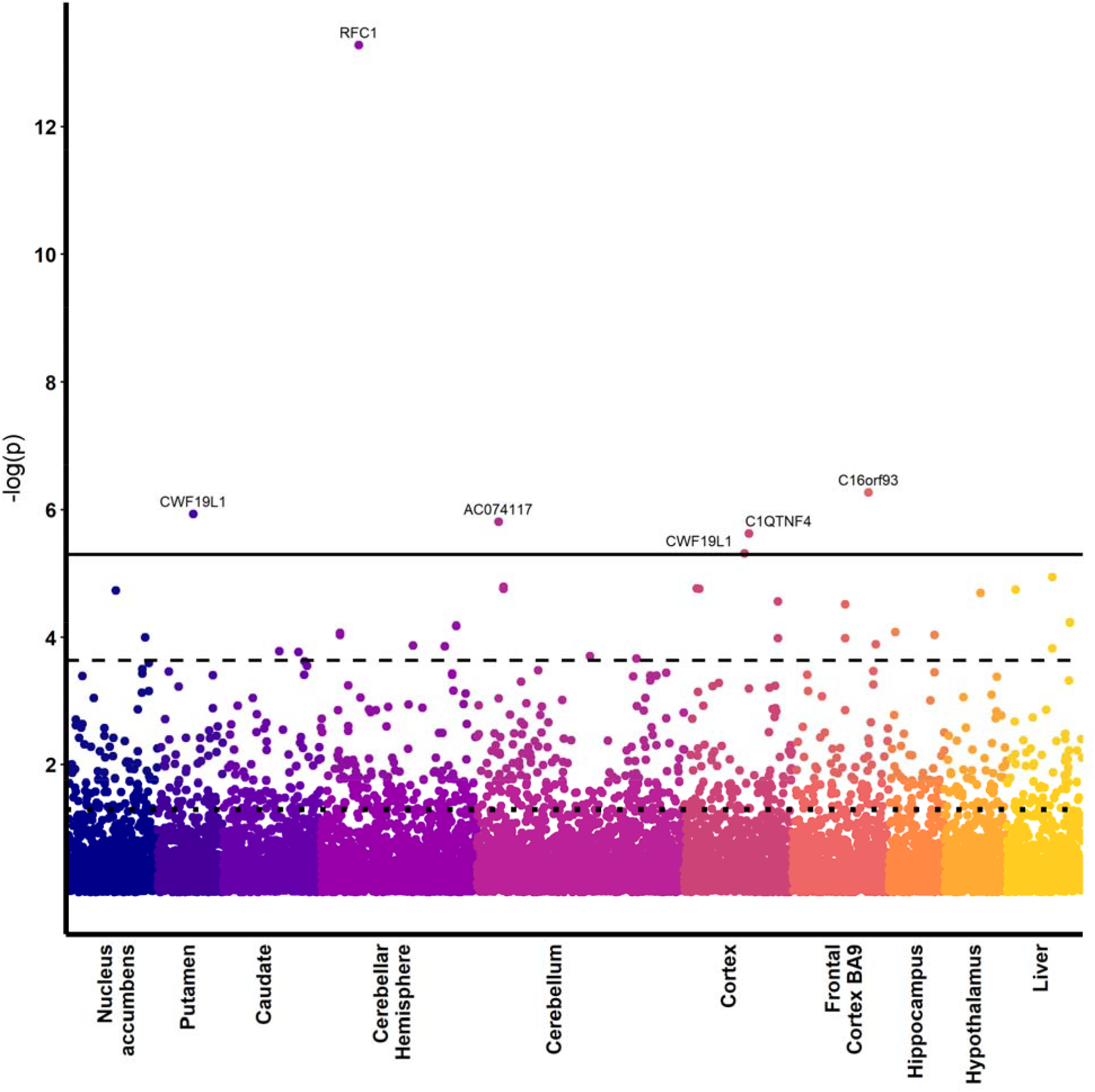
TWAS of alcohol consumption predicting gene expression. Genetic risk for alcohol consumption according to the UK Biobank GWAS (N=112,117) is associated with differences in human post-mortem gene expression (GTEx; Ns = 81 - 103), including frontal cortex BA9 (**Supplemental Data**). Notably, associations in the liver (far right) do not survive bonferroni-correction for multiple comparisons, though four are significant at a less-stringent FDR-based correction. Y-axis represents the significance of the association. Solid, dashed, and dotted lines represent Bonferroni-corrected, FDR-corrected, and nominally significant p-values, respectively.

**Table 2.**
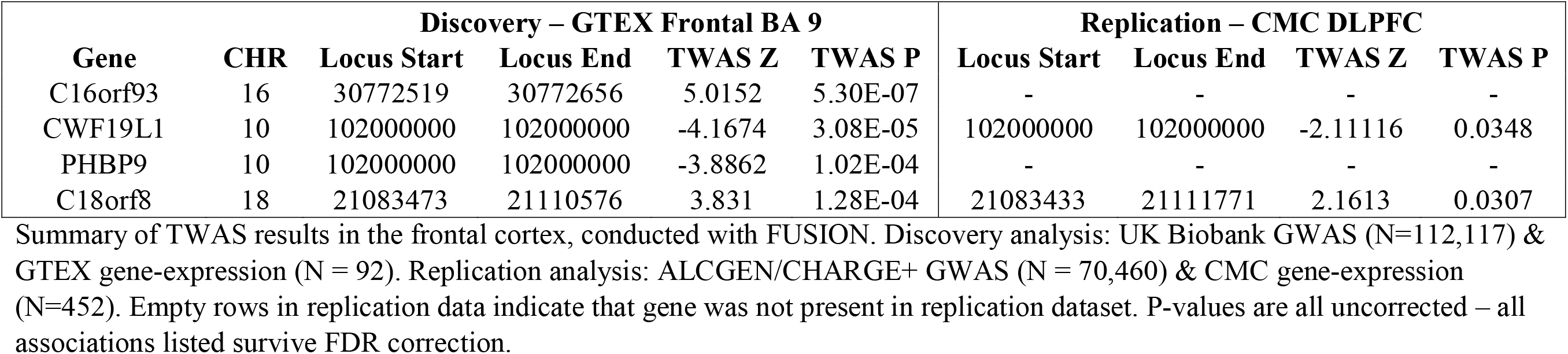
TWAS Discovery and Replication

| **Gene** | **CHR** | **Discovery – GTEX Frontal BA 9** |  |  |  | **Replication – CMC DLPFC** |  |  |  |
| --- | --- | --- | --- | --- | --- | --- | --- | --- | --- |
|  |  | **Locus Start** | **Locus End** | **TWAS Z** | **TWAS P** | **Locus Start** | **Locus End** | **TWAS Z** | **TWAS P** |
| C16orf93 | 16 | 30772519 | 30772656 | 5.0152 | 5.30E-07 | - | - | - | - |
| CWF19L1 | 10 | 102000000 | 102000000 | -4.1674 | 3.08E-05 | 102000000 | 102000000 | -2.11116 | 0.0348 |
| PHBP9 | 10 | 102000000 | 102000000 | -3.8862 | 1.02E-04 | - | - | - | - |
| C18orf8 | 18 | 21083473 | 21110576 | 3.831 | 1.28E-04 | 21083433 | 21111771 | 2.1613 | 0.0307 |

Summary of TWAS results in the frontal cortex, conducted with FUSION. Discovery analysis: UK Biobank GWAS (N=112,117) & GTEX gene-expression (N = 92). Replication analysis: ALCGEN/CHARGE+ GWAS (N = 70,460) & CMC gene-expression (N=452). Empty rows in replication data indicate that gene was not present in replication dataset. P-values are all uncorrected – all associations listed survive FDR correction.

An examination of putative behavioral mediators available in the DNS and HCP (i.e., executive function, delay discounting, self-reported impulsivity, negative urgency, and neuroticism) that might mediate links between brain structure and alcohol consumption were tested for association with GMV. Despite nominally significant associations between delayed-discounting and IQ, and GMV of the right frontal cortex in the DNS, none of these associations replicated within the HCP data or were robust to multiple testing correction (see **Supplement; Supplemental Tables 9-10**).

## DISCUSSION

We report convergent evidence that smaller right insula and DLPFC (i.e., middle and superior frontal gyri) GMVs plausibly represent genetically-conferred liabilities that promote early alcohol use. First, we show that smaller GMV of the right insula and DLPFC were replicably associated with alcohol use in two large neuroimaging samples. Second, family-based data provide evidence that these associations are attributable to shared genetic factors with no evidence that environmental factors contribute to this association. Third, reduced DLPFC volume prospectively predicted future alcohol use among young adults as well as alcohol use initiation during adolescence among children/adolescents who were unexposed to alcohol at baseline. Finally, we find replicable evidence that genomic risk for alcohol use is enriched among genes preferentially expressed within the DLPFC, and that genomic liability to alcohol use is associated with differential expression of *C16orf93*, *CWF19L1*, *C18orf8* in the DLPFC. Collectively, these convergent data suggest that lower GMV in middle/superior frontal gyri and insula may represent a preexisting genetic liability for drinking that could serve as a prognostic biomarker. Further, these data suggest the alcohol use in the general population does not induce reductions in GMV, at least as measured using MRI, as has been previously hypothesized (5, 8, 10). It is possible that reduced GMV in the superior and middle frontal gyri and insula may promote alcohol use, increasing the likelihood of heavy use, which may then further potentiate GMV loss in these regions and others (10, 15, 16).

A few notable points within our data require additional interpretation. First, in the longitudinal child/adolescent sample of baseline non-users (i.e., TAOS) we find that DLPFC GMV prospectively predicts an early age of drinking initiation. In the DNS longitudinal prospective data of young adults, reduced GMV in these regions also predicted future alcohol use, even after accounting for the extent of baseline alcohol use. However, this was only predictive up until age 20.85. It is possible that risk conferred by reduced GMV in the DLPFC is developmentally constrained and/or may be minimized by environmental differences in permissivity/legality, as the legal drinking age in the United States is 21 (65).

Second, we find no compelling evidence that behaviors speculated to contribute to alcohol use (e.g., executive function, negative urgency, and impulsivity) are associated with prefrontal or insula GMV, leaving the behavioral mechanisms through which these GMVs influence alcohol use unclear. That DLPFC GMV was negatively correlated with delay discounting and using alcohol to cope with stress in our young adult sample (DNS) at nominal levels of significance, while being unlinked in our young/middle aged adult sample (HCP; **Supplementary Tables 9&10**), is consistent with potentially developmentally constrained effects.

Third, genomic risk for alcohol use was enriched only within brain gene-sets. Brodmann Area 9, which overlaps with the DLPFC regions identified in our neuroimaging analyses was among the regions of strongest enrichment (Figure 4). Further, TWAS analyses revealed replicable evidence that genomic risk for alcohol use is associated with differential expression of *CWF19L1* and *C18orf8* within these regions. The function of these genes is not understood. Both genes have been previously implicated in psychopathology and related traits including schizophrenia, substance use, and cognition (**Supplementary Data**). Further, highlighting the potential importance of *CWF19L1* in brain development, rare mutations in *CWF19L1* are causes of autosomal recessive cerebellar ataxia (66, 67), which is characterized by a loss of control of bodily movements, as well as developmental delay and mental retardation.

Finally, given evidence that genetic liability is shared across substance use involvement (68) as well as other forms of psychopathology (69), our findings may generalize to other substances and generalized psychopathology risk. Genetically-informed and longitudinal studies enriched for other substance use and psychopathology, as well as large prospective studies, such as the Adolescent Brain Cognitive Development (ABCD) study could test this (70). While enrichment analyses implicate only brain pathways and TWAS identify replicable associations between genetic risk for alcohol consumption and gene expression in the frontal cortex, we cannot rule out the possibility that our observed effects are partially mediated by altered functioning of other pathways, such as alcohol metabolism in the liver (71).

While our study is limited by our sample size, particularly of discordant twins and longitudinal analyses, a major strength of our results is the convergent evidence provided by the different study designs (72). We note that our cross-sectional analyses of alcohol consumption and longitudinal analyses of adolescent use initiation, are the largest to-date that we know of. A primary limitation of our gene-expression analyses is that both of the gene-expression datasets included alcohol-exposed donors. Given the wide prevalence of alcohol use across the world (73), it will likely be impossible to ever definitively confirm in human adults that alcohol use is not confounding these results. Notably, none of the identified genes (*C16orf93*, *CWF19L1*, *C18orf8)* have been found to be differentially expressed in the frontal cortex of donors with alcoholism (74, 75). Our analyses are also limited by the omission of the insula from the gene-expression data, precluding a comparison of the gene-expression correlates between the insula and frontal cortex.

Limitations notwithstanding, our study provides convergent evidence that smaller GMV in the insula and DLPFC associated with alcohol use may represent a genetically-conferred liability that promotes early alcohol use. While this may in turn lead to accelerated volume loss within these and other regions, these findings challenge predominant interpretations that smaller brain volumes tied to alcohol use emerge primarily from the atrophy-inducing effects of alcohol. As larger prospective samples are acquired (e.g., ABCD), it will be interesting to test influential theories of substance use liability, and to examine the interplay of genetic risk and substance-use on the trajectories of brain development.

## Funding

Data for this study were provided by the Human Connectome Project, WU-Minn Consortium (principal investigators: David Van Essen, PhD, and Kamil Ugurbil, PhD; grant 1U54MH091657) funded by the 16 National Institutes of Health institutes and centers that support the National Institutes of Health Blueprint for Neuroscience Research, as well as by the McDonnell Center for Systems Neuroscience at Washington University. The Duke Neurogenetics Study is supported by Duke University and NIDA (DA033369). The Teen Alcohol Outcomes Study (TAOS) was supported by Duke University and NIAAA (AA016274) (DEW). DAAB was supported by NIH (T32-GM008151) and NSF (DGE-1143954). LF was supported by NIAAA (AA023693). CHD was supported by NIH (T32-DA007313 and T32-GM081739). AA was supported by NIDA (5K02DA32573). ARH receives additional support from NIDA (DA031579) and NIA (AG049789). LS was supported by K23 AA025399 and U01 DA041093. RB was supported by the Klingenstein Third Generation Research and NIH (R01-AG045231, R01-HD083614, R01-AG052564).

## Data Availability

All data pertaining to this study are available upon request to the corresponding author. Data sets can also be accessed at the locations listed:

Duke Neurogenetics Study (DNS): https://www.haririlab.com/projects/procedures.html

Human Connectome Project (HCP): https://www.humanconnectome.org/

Teen Alcohol Onset Study (TAOS): Requests for data access should be submitted to the study PI, Dr. Douglas Williamson -

UK Biobank: http://www.ukbiobank.ac.uk/

ALCGEN/CHARGE+: Requests for access to summary statistics should be submitted to the study PI, Dr. Gunter Schumann -

GTEx/CMC: Pre-computed gene expression weights were provided by the Gusev lab - http://gusevlab.org/projects/fusion/; https://gtexportal.org/; http://gusevlab.org/projects/fusion/; https://gtexportal.org/;

https://www.nimhgenetics.org/resources/commonmind

## Footnotes

1 Analyses repeated excluding non-drinkers (i.e., those who scored 0 on the AUDIT) yield equivalent results (see **Supplemental Figure 2**).

2 Notably, correcting for only tests within Brodmann Area 9 based on our neuroimaging results, CWF19L1 remains significant following Bonferroni correction.

