## Supplement for "Convergent evidence for predispositional effects of brain gray matter volume on alcohol consumption"

**Supplementary Information Table of Contents**

**Supplemental Methods:** Pages 3-14

**Supplemental Results:** Pages 15-18

**Supplemental Figures 1-5:** Pages 19 – 23

**Supplemental Tables 1-11:** Pages 24 - 35

**Supplemental References:** Pages 36-37

**Supplemental Methods**

**Duke Neurogenetics Study (DNS):** The DNS (cross-sectional; n=1334) assessed a wide range of behavioral, experiential, and biological phenotypes among young-adult (18-22 year-old) college students. Each participant provided informed written consent prior to participation in accord with the guidelines of the Duke University Medical Center Institutional Review Board and received $120 remuneration. All participants were in good general health and free of DNS exclusion criteria: (1) medical diagnosis of cancer, stroke, diabetes requiring insulin treatment, chronic kidney or liver disease or lifetime psychotic symptoms; (2) use of psychotropic, glucocorticoid or hypolipidemic medication, and (3) conditions affecting cerebral blood flow and metabolism (e.g., hypertension). DSM-IV Axis I and select Axis II disorders (Antisocial Personality Disorder and Borderline Personality Disorder) were assessed with the electronic Mini International Neuropsychiatric Interview (e-MINI) (1) and Structured Clinical Interview for the DSM-IV Axis II Personality Disorders (2). These disorders are not exclusionary as the DNS seeks to establish broad variability in multiple behavioral phenotypes related to psychopathology. Participants were excluded from analyses due to: 1) non-completion of T1 structural scans (n=10), 2) scanner-related artifacts in MRI data (n=12), 3) incidental structural abnormalities (n=4), 4) missing or incomplete data (n=4), and genetic anomalies (e.g., Kleinfelter’s syndrome; n=1). The final DNS sample consisted of 1,303 participants after quality assurance (**Supplementary Table 5**; age=19.70±1.25; 747 female; 258 with a DSM-IV Axis I disorder; **Supplementary Table 6**).

DNS participants were contacted every 3 months after initial study completion, and asked to complete a brief online assessment. Participants were entered into a lottery for a $50 gift-card following completion of each online assessment. Of the 734 participants who completed at least one online assessment (2,075 total responses), 705 completed the AUDIT questionnaire at least once, and 679 of these participants were among those included in initial DNS analyses (1,903 responses; **Supplemental Table 7**). Participants completed between 2 and 17 follow-ups (M=4.05, SD=2.62), between 28 and 1707 days after study completion (M=413.96, SD=331.22; age range: 18.33 - 23.82; **Supplementary Figure 5**).

**Human Connectome Project (HCP):** Data from participants contained in the HCP December 2015 public data release (N = 970), were considered for analyses. The HCP aims to recruit 1200 individuals (3-4 siblings per family, most including a twin pair) with the broad goal of examining individual differences in brain circuits and their relation to behavior and genetic background (3). Each participant provided informed written consent prior to participation in accord with the guidelines of the Washington University in St Louis Institutional Review Board and received $400 remuneration, as well as additional winnings ($5) and travel expenses. All participants were aged 22 to 35 years and free of the following exclusionary criteria: preterm birth, neurodevelopmental, neuropsychiatric, or neurologic disorders; a full list of exclusions is available in prior publication (4). Participants were excluded from analyses in the present study for poor quality structural MRI data (n=73), and non-completion of study questionnaires (n=3), resulting in a final sample of 894 (**Supplementary Table 5**; age=28.82±3.68; age range: 22-37; 393 males; 149 meeting criteria in a phone interview for a possible DSM-IV Axis I disorder (5) **Supplementary Table 6**).

**Teen Alcohol Outcomes Study (TAOS):** TAOS is a longitudinal study designed to examine the association between the development of depression and alcohol use disorders by recruiting adolescents aged 11 – 15 (N = 330) at high and low familial risk for depression (high: at least one first-degree and one second-degree relative with a lifetime history of major depression; low: no first-degree and minimal second-degree relatives (< 20%) with a lifetime history of depression). Participant diagnoses were assessed through structured clinical interviews with the adolescent and parent separately, using the Schedule for Affective Disorders and Schizophrenia for School-Age Children—Present and Lifetime Version (6). Participants were excluded if they met criteria for a substance use disorder or reported binge drinking at baseline (based on National Institute on Alcohol Abuse and Alcoholism guidelines). Participants were permitted to present with anxiety disorders in the high familial depression risk group (specific phobia, N=9; social phobia, N=6; panic disorder, N=1; generalized anxiety disorder, N=12). No other forms of psychopathology were present within the sample at baseline.

Participants were contacted every year to complete diagnostic interviews and questionnaires, and also underwent a follow-up MRI scanning session during the 2^nd^-4^th^ year of participation (these scans were not used in the present analysis). Participants provided assent, and parents provided written informed consent following procedures approved by the Institutional Review Board at the University of Texas Health Sciences Center at San Antonio, and received $165 remuneration, as well as additional winnings ($10), travel expenses, and $40 for each annual follow-up. Participants were excluded from analyses in the present study for non-completion of the MRI study session (n=17), poor quality structural MRI data (n=13), non-completion of follow-up visits (n=24), missing baseline self-report measures (n=40), and initiation of alcohol use prior to MRI scan (n=14), resulting in a final sample of n=223 (baseline age: 11-15; follow-up ages: 12-20; **Supplemental Table 5**).

**Alcohol Use Assessment**

**DNS:** Participants completed the 10-item Alcohol Use Disorders Identification Test (AUDIT), which was developed by the World Health Organization to screen for hazardous or dependent alcohol use patterns by assessing the frequency and nature of consumption over the past 12 months. The AUDIT had adequate internal consistency (α=0.81; M=5.22; SD=4.31; range 0-23). We computed the subscale score of the 3 items that correspond to the hazardous use or consumption domain of the AUDIT (AUDIT-C; Bush et al., 1998) (α=0.85; M=3.76; SD=2.64; range 0-12). Participants completed the AUDIT at baseline and during follow-up online assessments. Follow-up AUDIT questions were modified to ask about alcohol use following the participant’s last assessment.

**HCP:** Participants completed the Semi-Structured Assessment for the Genetics of Alcoholism (SSAGA) (5). From the SSAGA we created a metric, the modified lifetime AUDIT-C (mAUDIT-C), to approximate maximum lifetime AUDIT-C (α=0.786; M=3.42; SD=2.65; range 0-12). This used questions almost identical to those contained in the AUDIT-C, but with the difference that the SSAGA asks about frequency of drinking 5+ drinks in a 24-hour period, while the AUDIT asks about 6+ drinks.

**TAOS:** Adolescents were assessed at baseline and each annual follow-up session with the Substance Use Questionnaire (SUQ) (8). In addition to questions about other substances, the SUQ assesses lifetime exposure to alcohol (e.g., have you ever had a drink, have you ever been drunk, age of first drink) and onset of regular use (i.e., at least once per month for at least six months). The SUQ contains questions identical to those on the AUDIT-C. Thus, as in HCP analyses, a modified AUDIT-C score was created (mAUDIT-C: α=0.893; M=0.45; SD=1.26; range 0-9). Initiation of alcohol consumption was defined as attaining a score of 1 or greater on the mAUDIT-C (i.e. participant reports having consumed at least one full alcoholic beverage in the past year; N=82 initiated during the study; Age: M = 16.68, SD = 1.39, 14.12 – 19.64 yrs).

**Self-report Questionnaires and Behavioral Phenotypes Covariate Assessment**

**Socioeconomic Status – Income and Education**

**DNS:** Participants completed three Likert-scale questions, (scores range from 1-11) where higher scores reflect having a higher socioeconomic status (more money, education, and respected jobs). Participants were asked to place themselves, their biological father during their childhood and adolescence, and their biological mother during their childhood and adolescence, on this scale. These three responses were averaged to compute a proxy for the participant’s socioeconomic status (P-SES: α=0.816; M=7.44; SD=1.70; range 1.33-11). Since all participants in the DNS were enrolled in college at the time of the study, parental education (the average of the education of the participant’s female and male guardian) was used as a proxy, which ranged from 1 (some high school), to 9 (doctoral degree; MD, PhD, JD, or PharmD) (P-ED: α=0.70; M=7.09; SD=1.62).

**HCP:** Participants reported their total annual household income on a scale ranging from 1 (<$10,000), to 8 (>=$100,000) (SES: M=4.97; SD=2.18) as well as the number of years of education that they had completed (ED: M=14.89; SD=1.82; range=11-17).

**TAOS:** The parents of participants reported their annual income and their spouses annual income on a scale ranging from 0 (Less than $10,000) to 7 ($150,000 or more) (Self: M=4.27; SD=1.39; range=0-6. Spouse: M=4.12; SD=1.71; range=0-6.). Parents also reported their highest level of education and their spouses highest level of education on a 0 (Less than 9^th^ grade) to 6 (Graduate or Professional Degree) scale (Self: M=2.67; SD=12.36; range=0-9. Spouse: M=4.03; SD=2.06; range=0-9.). An overall estimate of SES was calculated by averaging the standardized values of these variables.

**Early-life stress**

**DNS:** Participants completed the 28-item Childhood Trauma Questionnaire (CTQ) (9), which asks participants to retrospectively report on the occurrence and frequency of emotional, physical, and sexual abuse as well as emotional and physical neglect before the age of 17 (α=0.88; M=33.55; SD=8.76; range 25-76). The instrument’s five subscales, each representing one type of abuse or neglect, have robust internal consistency and convergent validity with a clinician-rated interviews of childhood abuse (10). Total CTQ scores across the 5 subscales were used as a covariate.

**HCP:** The HCP did not include a measure of early-life stress.

**TAOS:** Participants completed the 28-item CTQ; total scores across the 5 subscales were used as a covariate in analyses (α=0.795; M=32.98; SD=8.76; range 25-64).

**Perceived Stress**

**DNS:** Participants completed the 10-item version of the Perceived Stress Scale (PSS) (11), which instructs participants to appraise how unpredictable, uncontrollable, and stressful their daily life was in the preceding week. The PSS had good internal consistency (α=0.86; M=14.66; SD=6.08; range 0-37).

**HCP:** HCP participants completed the same 10-item version of the PSS, which had good internal consistency (α=0.90; M=13.07; SD=5.76; range 0-35).

**TAOS:** TAOS participants completed the Stressful Life Events Schedule (SLES) (12), which assesses the presence of more than 80 possible stressors in the past 12 months, each rated on a 4-point scale (1-4). Each stressor is given a subjective stress score as rated by the adolescent and an objective stress rating by a consensus panel. For both subjective and objective stress, a summary score derived by summing the squares of each individual stressor. Herein we use the total subjective stress (M=31.52; SD=28.79; range 0-156).

**Additional Measures in TAOS**

**Depressive Symptoms:** As the TAOS sample was enriched for participants with a family history of depression, self-report of depressive symptoms was included as a covariate in all analyses. Participants completed the Mood and Feelings Questionnaire (MFQ) (13), a 33-item measure designed to detect clinically meaningful signs and symptoms of depressive disorders in children and adolescents (α=0.68; M=9.03; SD=8.01; range 0-52).

**Tanner Stage:** The tanner scale was used to assess pubertal status (Female: M=3.6, SD = 0.9; Male: M=2.97, SD=0.8) (14, 15).

**Variable transformation**

Sample demographics and comparisons, as well as associations of self-report measures with alcohol consumption, were computed in R (3.3.2) (16). Self-report questionnaire data were winsorized (to ± 3 SDs; DNS: AUDIT-C N=1, P-SES N=5, PSS N=6, CTQ N=25; HCP: aAUDIT-C N=3, PSS N=3; TAOS: SLES N=12, CTQ N=8; MFQC = 2) to maintain variability while limiting the influence of extreme outliers. Self-report variables with high skew (>1 or <-1) were transformed prior to analyses. Right skewed variables (DNS: CTQ skew=1.34; TAOS: CTQ skew = 1.16, SLES skew = 1.23) were log-transformed, while left-skewed variables (DNS: P-ED skew=-1.43) were squared.

**Magnetic Resonance Imaging Study Acquisition Parameters**

**DNS:** Two identical research-dedicated GE MR750 3 T scanners equipped with high-power high-duty-cycle 50-mT/m gradients at 200 T/m/s slew rate, and an eight-channel head coil for parallel imaging at high band-width up to 1 MHz were used to acquire data at the Duke-UNC Brain Imaging and Analysis Center (N=224, 17% of the sample, was scanned on the second scanner. Scanner is included as a covariate in all analyses). High-resolution T1-weighted images were obtained using a 3D Ax FSPGR BRAVO with the following parameters: TR = 8.148 s; TE = 3.22 ms; 162 sagittal slices; flip angle, 12°; FOV, 240 mm; matrix =256×256; slice thickness = 1 mm with no gap; and total scan time = 4 min and 13 s. Regional gray matter volumes were determined using the unified segmentation (17) and DARTEL normalization (18) modules in SPM12 (http://www.fil.ion.ucl.ac.uk/spm) (19). Using this approach, individual T1-weighted images were segmented into gray, white, and CSF images then non-linearly registered to the existing IXI template of 550 healthy subjects averaged in standard Montreal Neurological Institute space, available with VBM8 (http://dbm.neuro.uni-jena.de/vbm/). Subsequently, gray matter images were modulated for nonlinear effects of the high-dimensional normalization to preserve the total amount of signal from each region, and smoothed with an 8mm FWHM Gaussian kernel. The voxel size of processed images was 1.5×1.5×1.5 mm. A gray matter mask for subsequent analyses was created by thresholding the final stage (6th) IXI template at 0.1.

**HCP:** High-resolution (0.7-mm isotropic voxels) anatomical images were acquired using a customized Siemens Skyra 3-T scanner with a 32-channel head coil (20). Briefly, relevant steps for this study from the HCP processing pipeline within FSL v5.0.6 (21) included: (1) Gradient distortion correction, (2) Coregistration and averaging of T1 and T2 runs, (3) Linear registration of T1 and T2 runs, (4) FSL FNIRT brain extraction, (5) Field mad distortion correction, and (6) Bias field correction. Brain-extracted images were grey matter-segmented before being registered to the MNI 152 standard space using non-linear registration (22). The resulting images were averaged and flipped along the x-axis to create a left-right symmetric, study-specific grey matter template. Native grey matter images were then non-linearly registered to this study-specific template and multiplied by the Jacobian of the warp field to correct for local expansion (or contraction) due to the non-linear component of the spatial transformation. These images were then smoothed with an isotropic Gaussian kernel with a sigma of 4 mm.

**TAOS:** Imaging data was collected using a Siemens 3T Trio scanner located at the Research Imaging Institute at the University of Texas Health Science Center, San Antonio (UTHSCSA). The study used an MRI protocol specifically optimized for GM thickness measurement (23). The protocol was designed to collect data to resolve the cortical ribbon across to cortex using isotropic spatial resolution of 0.8mm, voxel size =0.5mm^.^ T1-weighted contrast was achieved using a magnetization prepared sequence with an adiabatic inversion contrast-forming pulse (scan parameters: TE/TR/TI=3.04/2100/785 ms, flip angle=11 degrees). The processing of T1-weighted images consisted of removing non-brain tissues, global spatial normalization and radio frequency (RF) inhomogeneity correction. Non-brain tissues such as skin, muscle and fat was removed using an automated skull stripping procedure and images were corrected for radio-frequency (RF) inhomogeneity (24). A retrospective motion-correction technique was used to reduce subject motion-related artifacts (25). Next, images were imported into the structural analysis package, BrainVisa, and processed using its cortical extraction and parcellation pipelines (25). This pipeline extracts the pial and grey matter and white matter (WM) interface surfaces, performs extraction, labeling and verification of sulcal surfaces (26) and segments the cortical landscape into 15 cortical regions using the primary sulcal structures.

**Post-hoc Associations of volume with behavior**

**Methods**

Following the observation that frontal and insula volume are predictive of future alcohol use and initiation, are predispositional to alcohol consumption, and that genetic risk for alcohol consumption is associated with changes in gene expression in the frontal cortex (**Results**), *post-hoc* exploratory analyses sought to test whether the effects of volume on alcohol consumption are mediated by cognitive or behavioral measures. Associations between total volume of the ROI, as well as volume of each significant cluster, and three classes of outcomes were tested: impulsivity, negative urgency, and intelligence. As measures of negative urgency were not collected in the HCP, associations with neuroticism were additionally examined. The association between impulsivity and alcohol consumption is well established (27, 28), and measures of impulsivity have been shown to be associated with the structure of both the insula and frontal cortex in large samples (29–32). Negative urgency, a facet of impulsivity characterized by risky decision making when one is experiencing negative emotions, is associated with problematic drinking (33–35), and shows correlations with neural responses to reward (36), as well as with structure of the frontal cortex (37). Childhood intelligence has been observed to be predictive of adult alcohol consumption (38), with evidence of genetic correlations between proxy-measures of IQ (i.e. educational attainment) and alcohol consumption (39), and brain structure has been repeatedly linked to intelligence, with evidence that the two share genetic underpinnings (40–42).

**Delay Discounting**

In delay discounting tasks participants choose between hypothetical amounts of money available immediately or after a delay. By varying the amount of money available immediately and the number of days that one would have to wait for the delayed money, participant ‘indifference points’ can be identified wherein the participant is equally likely to choose a smaller reward sooner versus a larger reward later. A preference for a smaller reward sooner (i.e., delay discounting) is considered a behavioral index of impulsivity (43). The DNS and HCP used different protocols which are described below.

**DNS:** All combinations of immediate reward (varying from $0.10 to $105) and delay intervals [0, 7, 30, 90, 180, 365, or 1825 (i.e., 5 years) days] for the delayed reward of $100 were presented on a computer screen in randomized order (44, 45). 146 participants did not complete the delay discounting task, resulting in a final DNS sample of 1,157 for delay discounting analyses.

**HCP:** The delayed-reward amount was set to $200, with delays of 1, 6, 12 (1 year), 36 (3 years), 60 (5 years), or 120 months (10 years) presented in the order: 6, 36, 1, 60, 120, 12 months (46). Immediate reward amounts were adjusted on a trial-by-trial basis based upon participant response. Six participants did not complete the delayed-discounting task.

**Impulsivity**

DNS participants completed the 30-item self-report Barratt Impulsiveness Scale (BIS) (47). HCP participants completed the Achenbach Adult Self-Report (ASR) for Ages 18-59 (48). As in prior reports (49), a coarse measure of impulsivity (ASR-Imp) was computed from three questions in the ADHD subscale of the ASR. **DNS**: Barratt Impulsiveness Scale assesses the personality/behavioral construct of impulsiveness and had good internal consistency (α=0.84; M=61.69; SD=9.55; range 37-113). **HCP**: Our Achenbach Adult Self-Report (ASR) impulsivity composite had acceptable internal consistency (ASR-Imp; α=0.79; M=1.29; SD=1.24; range 0-6).

**Neuroticism**

Participants in the DNS completed the 240-item NEO Personality Inventory-Revised (50) (Neuroticism: α=0.85; M=86.04; SD=22.65; range: 37-113). Six participants did not complete the NEO. Participants in the HCP completed the 60-item NEO Five-Factor Inventory (50) (Neuroticism: α=0.83; M=16.43; SD=7.17; range: 0 - 43).

**Negative Urgency**

Participants completed two measures of negative urgency – the Impulsivity sub-scale of the NEO-PI-R (50) (α=0.71; M=17.07; SD=4.60; range: 3-32), and the substance-sue subscale of the brief COPE (BCOPE-sub) inventory. The BCOPE-sub consists of two items that assess how frequently respondents use drugs and alcohol as a coping mechanism (51) (α = 0.92; M=2.53; SD=1.11; range: 0-8).

**Intelligence**

**DNS:** Intelligence was assessed using the Wechsler Abbreviated Scale of Intelligence Second Edition (WASI-II) 2-subtest version (52), consisting of the Vocabulary and Matrix Reasoning subtests. The total score was computed as the sum of age-adjusted performance on the two sub-tests. 28 participants did not complete the WASI-II. **HCP:** Intelligence was assessed using the NIH toolbox (53). Fluid intelligence was assessed using the Flanker Inhibitory Control and Attention Test, Picture Sequence Memory Test, List Sorting Test, Pattern Comparison Test, and Dimensional Change Card Sort Test. Crystallized intelligence was assessed using the Oral Reading and Picture Vocabulary tests. Age-adjusted scores were averaged for fluid and crystallized intelligence, respectively, which were then averaged to form a measure of intelligence (54).

**Statistical Analysis**

Analyses were conducted in R (3.4.2) (16). Covariates were identical to those used in neuroimaging analyses. As in other analyses, variables were transformed to correct for skew, and were winsorized as needed. All variables were z-scored. Linear regression models were used to test whether extracted brain volume predicted behavioral and self-report outcomes. Analyses in the HCP used linear mixed-effects models (55), which controlled for family as a random effect, and sibling-status (i.e. DZ twin, MZ twin, half-sibling) was entered as fixed-effect covariates. FDR-correction for multiple comparisons was applied the entire set of analyses for each study.

**Figure Creation**

Figures 2 &3, and Supplemental Figures 1-3 were made in R (3.4.2) (R Core Team, 2014). The packages ‘ggplot2’ (56) and ‘ggpubr’ (57) were used to generate the figures, and the package ‘viridis’ (58) was used for the color-palette. ‘ggrepel’ (59) was used in Supplemental Figure 2 for gene-labels.

**Supplemental Results**

**Comparison of discovery and replication samples**

Sample comparisons are presented in **Supplemental Table 5**. The samples differed by sex, which was driven by fewer female participants in the TAOS sample (the DNS and HCP do not differ). Consistent with the recruitment of non-overlapping aged samples, HCP participants were all older than DNS participants, who were all older than TAOS participants. A significantly larger proportion of the participants from the HCP were Caucasian or African/African-American, while the DNS had a larger portion of Asian/Asian-American and Multi-racial/Native-American/Other participants. TAOS had the largest proportion of Hispanic participants. DNS participants had significantly higher AUDIT-C scores (**Supplemental Figure 1**), which is unsurprising given observations of elevated hazardous alcohol consumption among younger college-aged populations (60). Similarly, the higher levels of self-reported perceived stress among DNS participants is consistent with prior observations of increased stress in college-student samples (61).

**Association of covariates with alcohol consumption**

Association of covariates with alcohol consumption are presented in **Supplemental Table 8**. In the DNS and HCP men reported higher levels of alcohol consumption than women, consistent with prior reports of sex-differences in alcohol use (62). Notably, in TAOS, sex did not differ between initiators and non-initiators. Presence of a non-substance-related diagnosis was not associated with alcohol use in either the DNS or HCP. Consistent with prior reports, participants of European descent reported increased levels, while participants of African descent reported decreased levels (63, 64). In the DNS participants of Asian descent and of Multi-racial or Native-American descent reported the lowest amount of alcohol consumption (64). These associations did not reach significance in the HCP, likely due to the smaller number of participants in these groups in this sample. Age was associated with alcohol consumption in all samples, though in differing directions, which is consistent with observations in North American samples that alcohol consumption increases once young adults turn 21 (legal drinking age), and subsequently decreases as participants age (62, 65). Of note, in TAOS alcohol-consumption initiators were younger at baseline. In the DNS, perceived SES (P-SES) and parental education (P-ED) were both positively correlated with alcohol consumption, while no association was observed with these phenotypes in the HCP or in TAOS. Perceived stress was not associated with alcohol consumption in any sample, which is consistent with some prior reports (66, 67). Childhood trauma was associated with alcohol consumption in the DNS, and was trending in TAOS, consistent with an extensive literature indicating that early trauma increases risk for substance use (68).

**Longitudinal Data Results**

DNS: Differences between longitudinal responders and non-responders are presented in **Supplemental Table 7**. Participants who completed at least one follow-up questionnaire were younger, had lower AUDIT-C and total AUDIT scores. Further, responders were more likely to be female and white and less likely to be black. Of the four brain volume x age interactions modeled (Frontal x Linear-Age, Frontal x Quadratic-Age, etc.), only the interaction between right middle/superior frontal volume and the linear change in age was significant after fdr-correction for multiple comparisons (β=0.151, t=3.976, p=0.0007, p-fdr=0.008; **Figure 3, Supplemental Table 3**). Examination of regions of significance found that lower frontal volume predicted greater drinking before the age of 20.85 years.

**TAOS**: Of the n=223 adolescents who were non-drinkers at baseline that were included in analyses, n=82 (36%) reported having consumed at least one full alcoholic beverage during one of their follow-up interviews (i.e. they initiated consumption). Of the four brain volume x age interactions modeled (Mid-Frontal x Linear-Age, Mid-Frontal x Quadratic-Age, etc.), the interactions between both right middle and superior frontal volume and the linear change in age were significant after fdr-correction for multiple comparisons (Middle frontal: β=-57.04, t= -2.37, p-fdr =	0.036; Superior frontal: β=-60.74, t = -2.43, p-fdr=0.036; **Figure 3, Supplemental Table 4**).

**Associations of brain volume with behavior**

Extracted volume of the frontal and insula ROI, and volume of each significant cluster, were not significantly associated with any of the behavioral measures, in either sample, after correcting for multiple testing (**Supplemental Tables 9&10**). This suggests that negative urgency, impulsivity, or IQ do not fully mediate the association between brain volume and alcohol consumption. Given nominally significant results for some of these analyses within our DNS sample, it is possible that different behavioral mechanisms might represent risk at different ages and that these null results may represent type II error that larger samples might detect.

**Supplemental Figure 1: Comparison of AUDIT-C distribution in HCP and DNS**

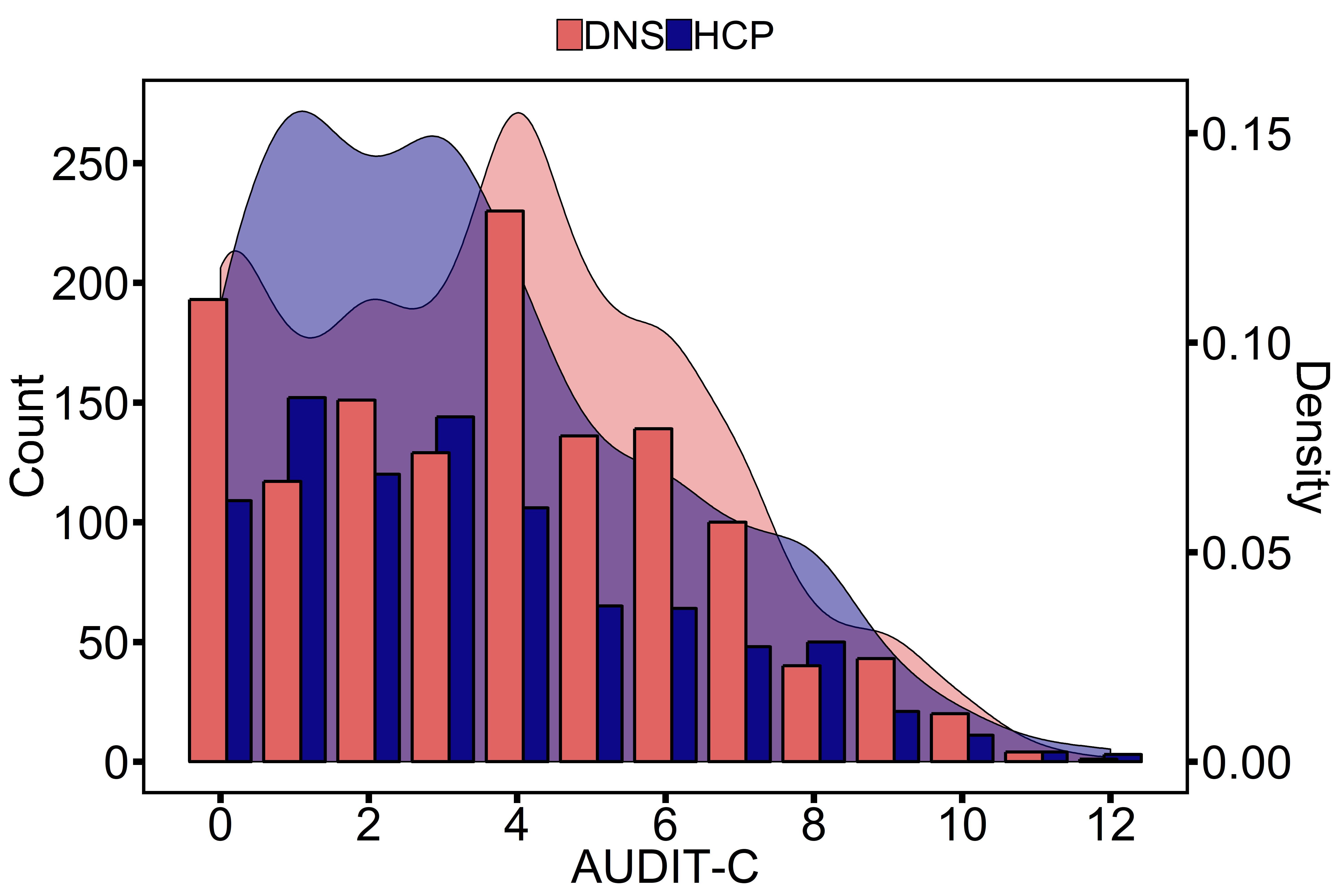

Comparison of distribution of AUDIT-C scores in the DNS (pink) and HCP (blue; aAUDIT-C).

**Supplemental Figure 2: Comparison of effects in the DNS and HCP**

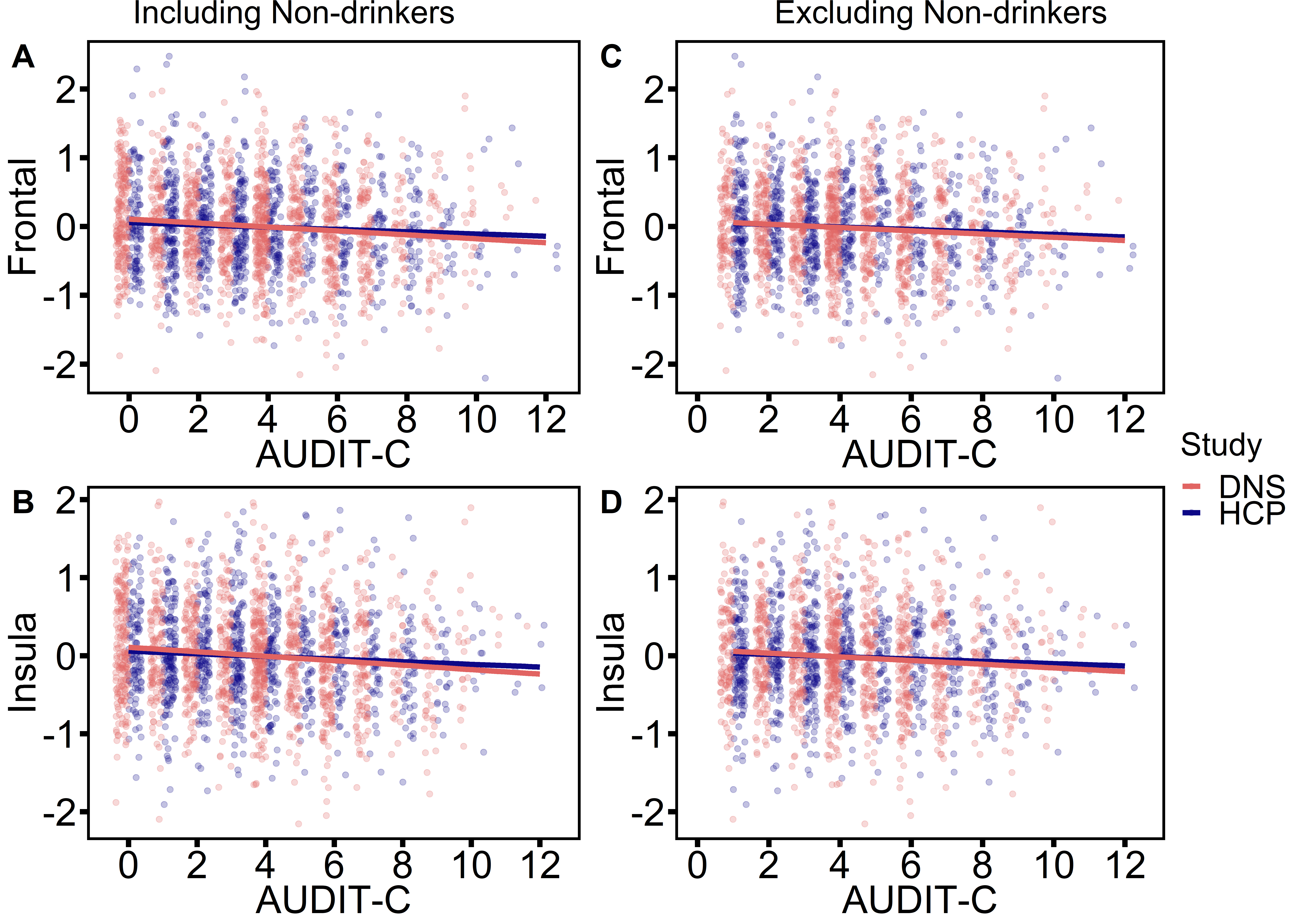

Comparisons of effects in the DNS (pink) and HCP (blue) between extracted volume of entire aal-atlas defined ROIs for the right middle and superior frontal cortex (‘Frontal’, **A** & **C**) and right insula (‘Insula’, **B** & **D**), and AUDIT-C scores. Brain-volume was standardized, and is corrected for all covariates used in models (see **Methods**), which were all standardized (z-scored). Results are presented including non-drinkers (AUDIT-C = 0, **A**&**B**) and excluding non-drinkers (AUDIT-C > 0, **C**&**D**). The association statistics are: **A)** DNS: β= -0.087, se=0.019 t= -4.46, p= 8.8x10^-6^; HCP: β= -0.077, se=0.033 t= -2.3, p= 0.022. **B)** DNS: β= -0.077, se=0.019 t= -3.97, p= 7.5x10^-5^; HCP: β= -0.082, se=0.032 t= -2.57, p= 0.010. **C)** DNS: β= -0.062, se=0.022 t= -2.87, p= 0.004; HCP: β= -0.082, se=0.035 t= -2.3, p= 0.019. **D)** DNS: β= -0.066, se=0.021 t= -3.07, p= 0.002; HCP: β= -0.080, se=0.033 t= -2.27, p= 0.018.

**Supplemental Figure 3: Overlap of HCP and DNS clusters at an uncorrected threshold**

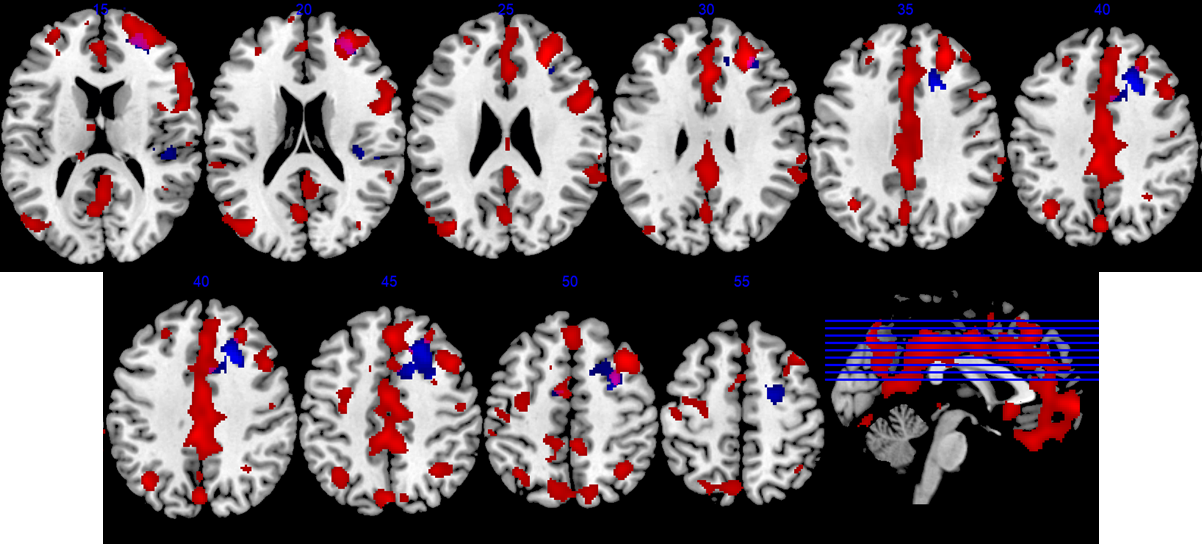

Overlap of volumetric associations in the frontal cortex with alcohol consumption in the DNS (red) and HCP (blue) samples (purple=overlap). Associations are displayed at an uncorrected threshold (DNS: p<0.001; HCP: p<0.05). For display purposes, different statistical thresholds were used due to the differential power of the two studies.

**Supplemental Figure 4: Models tested in discordant sibling analyses**

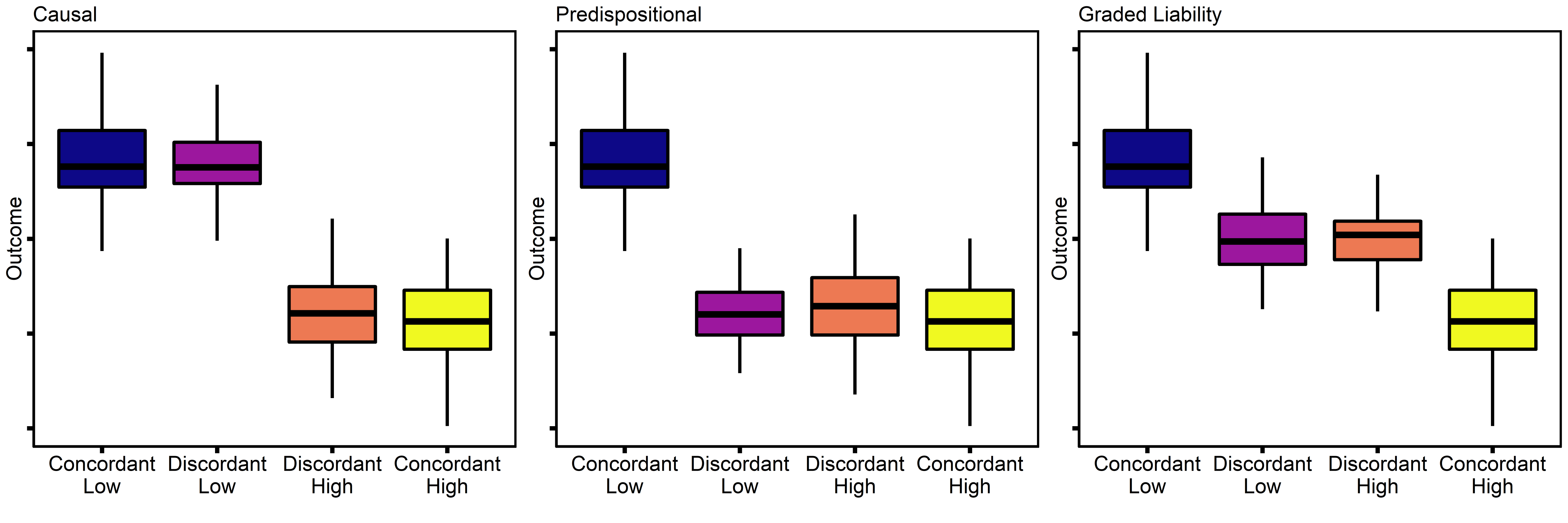

Schematics representing hypothetical models tested with contrast-codes in HCP discordant sibling analyses. **Causal:** The causal contrast tests whether siblings who had high levels of exposure (i.e. higher scores on the aAUDIT-C) differed from siblings with low exposure, on the outcome measure (i.e. brain volume). In particular, the model tests whether siblings who are discordant, that is who are at opposite ends of the distribution on the exposure, also differ in the outcome. As siblings share genetics and environment, a difference is indicative of a putatively causal effect of the exposure on the outcome. **Predispositional:** The predispositional contrast tests whether any amount of genetic-risk for the exposure, regardless of actual exposure, produces a similar outcome. This contrast suggests that the genetics shared between the low and high discordant siblings are sufficient for the low-discordant siblings to have an outcome similar to siblings that are both concordant for high-exposure. **Graded Liability:** The graded liability contrast tests (1) whether discordant siblings are similar to each other and (2) whether discordant pairs are intermediate between pairs that are concordant for low and high-exposure, respectively. The discordant siblings with high exposure differ from concordant siblings with high exposure, despite having the same level of actual exposure. This indicates that the degree of genetic risk matters for the outcome, as that the discordant siblings have lower genetic risk.

**Supplemental Figure 5: Distribution of responses to DNS follow-up questionnaire**

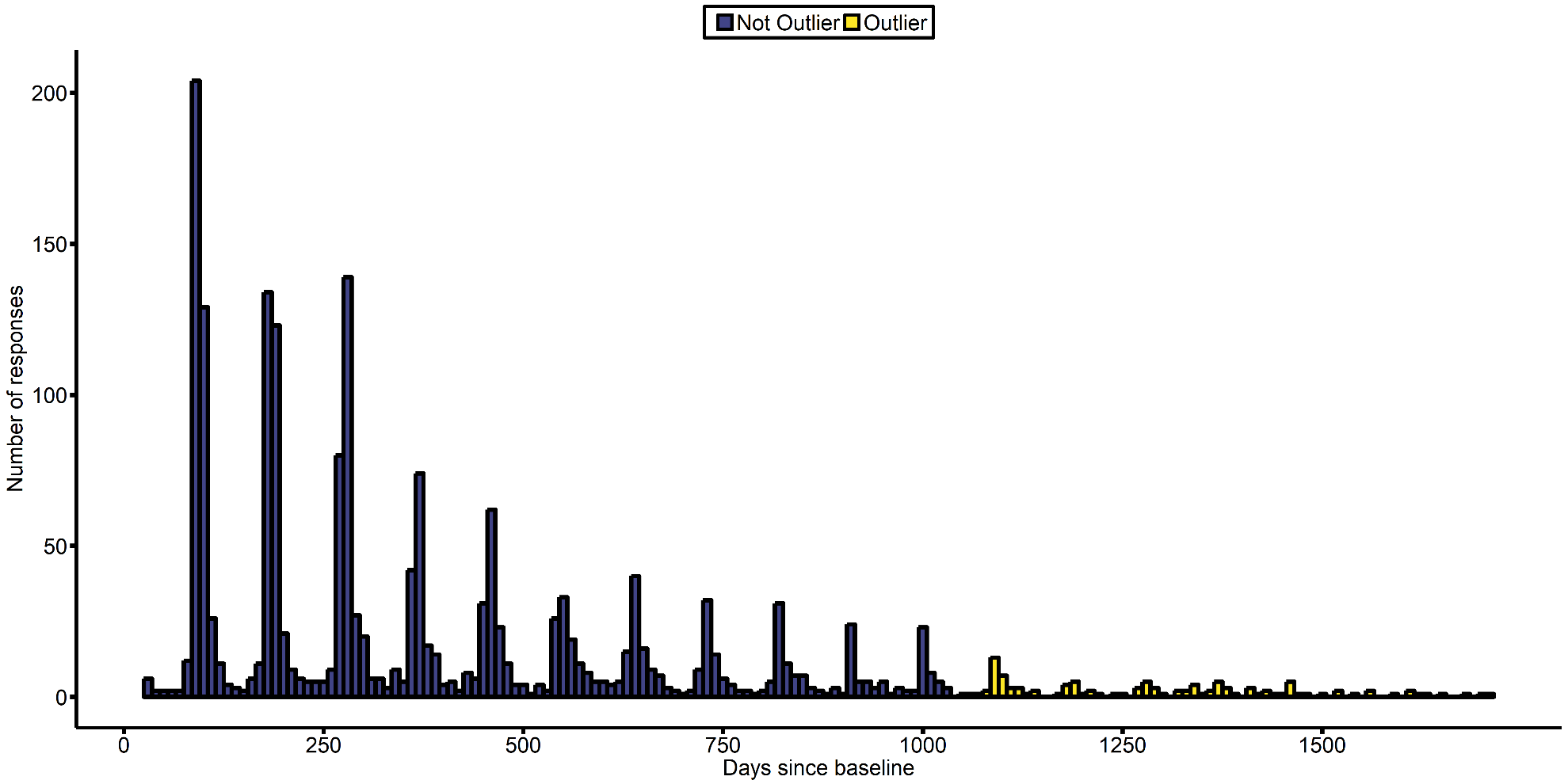

Histogram of responses to the DNS online follow-up questionnaire, which was emailed to participants every 3 months. Color indicates whether the clustering algorithm indicated the response to be an outlier or not – outliers occurred > 1053 days after the baseline visit and were excluded from analyses. Bins have a width of 10 days.

**Supplemental Table 1. Heritability and genetic correlation between gray-matter volume and alcohol consumption in the HCP**

|  | **mAUDIT-C** | **Right Insula** | **Right Middle/Superior Frontal Cortex** |
| --- | --- | --- | --- |
| **h^2^ (SE)** | 0.5179 (0.0541) | 0.6883 (0.0392) | 0.7746 (0.0307) |
| ***p*** | **5.25x10^-18^** | **1.97x10^-32^** | **3.62x10^-40^** |
| **ρ_p_** | - | -0.1349 | -0.114 |
| ***p*** | - | **0.0006** | **0.0033** |
| **ρ_g_ (SE)** | - | -0.2314 (0.076) | -0.2192 (0.0784) |
| ***p*** | - | **0.0022** | **0.0054** |
| **ρ_e_ (SE)** | - | 0.0294 (0.0825) | 0.0483 (0.0787) |
| ***p*** | - | 0.7214 | 0.541 |

h^2^ = heritability; ρ_p_ = phenotypic correlation; ρ_g_ = genetic correlation; ρ_e_ = environmental correlation

**Supplemental Table 2. Discordant sibling analysis in the HCP**

| **Region** | **Variable** | **Estimate** | **t** | **p** |
| --- | --- | --- | --- | --- |
| **Right Insula** | **ICV** | -5.78x10^-8^ | -4.629 | **4.78x10^-6^** |
|  | **Sex** | 0.0052 | 0.892 | 0.3733 |
|  | **Age** | -0.0030 | -5.571 | **4.27x10^-8^** |
|  | **MZ** | -0.0057 | -1.596 | 0.1111 |
|  | **DZ** | -0.0124 | -2.659 | **0.0081** |
|  | **W** | 0.0179 | 2.041 | **0.0419** |
|  | **B** | -0.0026 | -0.253 | 0.8007 |
|  | **A** | 0.0066 | 0.525 | 0.5998 |
|  | **SES** | 0.0002 | 0.289 | 0.7726 |
|  | **ED** | 1.20x10^-5^ | 0.362 | 0.7176 |
|  | **DX – Non Substance** | -0.0054 | -1.29 | 0.1978 |
|  | **PSS** | 0.0002 | 0.967 | 0.3343 |
|  | **Causal** | 0.0009 | 0.412 | 0.6806 |
|  | **Graded** | -0.0037 | -1.974 | **0.0491** |
|  | **Predispositonal** | 0.0037 | 3.479 | **0.0006** |
| **Region** | **Variable** | **Estimate** | **t** | **p** |
| **Right Middle/Superior Frontal Cortex** | **ICV** | 1.38x10^-8^ | 1.315 | 0.1891 |
|  | **Sex** | 0.0124 | 2.584 | **0.0103** |
|  | **Age** | -0.0029 | -6.438 | **3.01x10^-10^** |
|  | **MZ** | -4.89x10^-5^ | -0.016 | 0.9871 |
|  | **DZ** | -0.0059 | -1.505 | 0.1329 |
|  | **W** | 0.0139 | 1.899 | 0.0583 |
|  | **B** | 0.0208 | 2.387 | **0.0175** |
|  | **A** | 0.0217 | 2.058 | **0.0402** |
|  | **SES** | 0.0008 | 1.06 | 0.2896 |
|  | **ED** | -1.87x10^-5^ | -0.665 | 0.5067 |
|  | **DX – Non Substance** | 0.0092 | 2.557 | **0.0109** |
|  | **PSS** | 0.0002 | 1.042 | 0.2980 |
|  | **Causal** | -0.0020 | -1.037 | 0.3006 |
|  | **Graded** | 0.0006 | 0.411 | 0.6810 |
|  | **Predispositonal** | 0.0020 | 2.193 | **0.0290** |

SES = Socioeconomic status, PSS = Perceived Stress Scale, ED = education

**Supplemental Table 3. Regression analyses of the association between brain volume and longitudinal alcohol consumption in DNS**

| Variable | **β** | **Std.Error** | **DF** | **t-value** | **p-value** |
| --- | --- | --- | --- | --- | --- |
| (Intercept) | **0.114** | **0.056** | **1726** | **2.022** | **0.043** |
| Age-linear | **0.085** | **0.037** | **1726** | **2.266** | **0.024** |
| Age-quadratic | -0.019 | 0.031 | 1726 | -0.618 | 0.537 |
| Baseline Age | -0.026 | 0.048 | 646 | -0.546 | 0.585 |
| Sex | **-0.164** | **0.044** | **646** | **-3.698** | **2.35x10^-4^** |
| W | **0.161** | **0.062** | **646** | **2.615** | **0.009** |
| B | 0.032 | 0.045 | 646 | 0.714 | 0.476 |
| A | -0.047 | 0.058 | 646 | -0.809 | 0.419 |
| H | **0.093** | **0.039** | **646** | **2.360** | **0.019** |
| DX - Non Substance | 0.016 | 0.035 | 646 | 0.440 | 0.660 |
| PSS | -0.039 | 0.038 | 646 | -1.025 | 0.306 |
| P-SES | 0.066 | 0.041 | 646 | 1.594 | 0.111 |
| CTQ | 0.014 | 0.038 | 646 | 0.375 | 0.708 |
| P-ED | 0.074 | 0.040 | 646 | 1.845 | 0.065 |
| Scanner | 0.029 | 0.032 | 646 | 0.932 | 0.351 |
| ICV | 0.107 | 0.059 | 646 | 1.795 | 0.073 |
| Frontal | **-0.115** | **0.053** | **646** | **-2.185** | **0.029** |
| Age-linear x Frontal | **0.151** | **0.038** | **1726** | **3.976** | **7.29x10^-5^** |
| Age-quadratic x Frontal | 0.007 | 0.025 | 1726 | 0.275 | 0.783 |
| Age-quadratic x Baseline Age | **-0.046** | **0.018** | **1726** | **-2.490** | **0.013** |
| Age-quadratic x Sex | **-0.057** | **0.022** | **1726** | **-2.602** | **0.009** |
| Age-quadratic x W | -0.001 | 0.031 | 1726 | -0.022 | 0.982 |
| Age-quadratic x B | -0.009 | 0.022 | 1726 | -0.411 | 0.681 |
| Age-quadratic x A | 0.009 | 0.028 | 1726 | 0.322 | 0.747 |
| Age-quadratic x H | -0.009 | 0.021 | 1726 | -0.413 | 0.679 |
| Age-quadratic x DX - Non Substance | 0.017 | 0.018 | 1726 | 0.953 | 0.341 |
| Age-quadratic x PSS | 0.023 | 0.018 | 1726 | 1.269 | 0.205 |
| Age-quadratic x P-SES | 0.019 | 0.021 | 1726 | 0.883 | 0.377 |
| Age-quadratic x CTQ | 0.015 | 0.018 | 1726 | 0.799 | 0.424 |
| Age-quadratic x P-ED | -0.006 | 0.020 | 1726 | -0.284 | 0.777 |
| Age-quadratic x Scanner | 0.009 | 0.018 | 1726 | 0.479 | 0.632 |
| Age-quadratic x ICV | -0.026 | 0.028 | 1726 | -0.936 | 0.349 |
| Age-linear x Baseline Age | -0.019 | 0.048 | 1726 | -0.390 | 0.696 |
| Age-linear x Sex | 0.057 | 0.031 | 1726 | 1.876 | 0.061 |
| Age-linear x W | -0.014 | 0.042 | 1726 | -0.325 | 0.745 |
| Age-linear x B | -0.021 | 0.030 | 1726 | -0.702 | 0.483 |
| Age-linear x A | -0.010 | 0.039 | 1726 | -0.256 | 0.798 |
| Age-linear x H | 0.042 | 0.028 | 1726 | 1.519 | 0.129 |
| Age-linear x DX - Non Substance | **-0.053** | **0.025** | **1726** | **-2.153** | **0.031** |
| Age-linear x PSS | -0.015 | 0.025 | 1726 | -0.595 | 0.552 |
| Age-linear x P-SES | -0.047 | 0.028 | 1726 | -1.654 | 0.098 |
| Age-linear x CTQ | 0.004 | 0.026 | 1726 | 0.160 | 0.873 |
| Age-linear x P-ED | 0.024 | 0.027 | 1726 | 0.889 | 0.374 |
| Age-linear x Scanner | 0.018 | 0.024 | 1726 | 0.764 | 0.445 |
| Age-linear x ICV | **-0.090** | **0.040** | **1726** | **-2.243** | **0.025** |
| Baseline Age x Frontal | -0.016 | 0.041 | 646 | -0.386 | 0.700 |
| Sex x Frontal | 0.073 | 0.051 | 646 | 1.442 | 0.150 |
| W x Frontal | -0.043 | 0.065 | 646 | -0.669 | 0.503 |
| B x Frontal | 0.035 | 0.046 | 646 | 0.758 | 0.449 |
| A x Frontal | -0.056 | 0.060 | 646 | -0.924 | 0.356 |
| H x Frontal | -0.018 | 0.043 | 646 | -0.429 | 0.668 |
| DX - Non Substance x Frontal | 0.011 | 0.034 | 646 | 0.322 | 0.748 |
| PSS x Frontal | 0.023 | 0.038 | 646 | 0.605 | 0.546 |
| P-SES x Frontal | 0.009 | 0.043 | 646 | 0.213 | 0.832 |
| CTQ x Frontal | -0.052 | 0.037 | 646 | -1.407 | 0.160 |
| P-ED x Frontal | 0.034 | 0.042 | 646 | 0.819 | 0.413 |
| Scanner x Frontal | 0.001 | 0.035 | 646 | 0.038 | 0.970 |
| ICV x Frontal | 0.008 | 0.044 | 646 | 0.195 | 0.846 |

Volume of the right middle/superior frontal cortex is associated with future change in alcohol consumption (p-fdr=4x10^-4^). Standardized effects and uncorrected p-values are presented.

**Supplemental Table 4. Regression analyses of the association between brain volume and future alcohol use initiation in TAOS.**

|  | **Middle** | **Frontal** |  |  | **Superior** | **Frontal** |  |  |
| --- | --- | --- | --- | --- | --- | --- | --- | --- |
|  | **β** | **SE** | **z** | **p** | **β** | **SE** | **z** | **p** |
| **(Intercept)** | -6.64 | 156.30 | -0.04 | 0.97 | -6.35 | 132.19 | -0.05 | 0.96 |
| **Age-linear** | 170.77 | 5767.82 | 0.03 | 0.98 | 157.94 | 2198.72 | 0.07 | 0.94 |
| **Age-quadratic** | -27.21 | 4154.01 | -0.01 | 0.99 | -25.34 | 1750.80 | -0.01 | 0.99 |
| **Sex** | -0.04 | 0.94 | -0.04 | 0.97 | 0.15 | 0.80 | 0.19 | 0.85 |
| **W** | 0.99 | 1.68 | 0.59 | 0.56 | 0.69 | 1.38 | 0.50 | 0.62 |
| **H** | -0.18 | 1.63 | -0.11 | 0.91 | -0.23 | 1.32 | -0.17 | 0.86 |
| **High-risk** | 0.30 | 0.82 | 0.37 | 0.71 | 0.05 | 0.73 | 0.07 | 0.95 |
| **GenPop** | -2.70 | 1217.35 | 0.00 | 1.00 | -3.35 | 1029.61 | 0.00 | 1.00 |
| **SLES** | 0.83 | 0.93 | 0.89 | 0.37 | 0.94 | 0.78 | 1.20 | 0.23 |
| **SES** | -0.10 | 0.83 | -0.13 | 0.90 | -0.34 | 0.70 | -0.48 | 0.63 |
| **CTQ** | 1.15 | 0.77 | 1.50 | 0.13 | 1.02 | 0.68 | 1.51 | 0.13 |
| **Tanner** | -0.51 | 0.83 | -0.61 | 0.54 | -0.28 | 0.73 | -0.39 | 0.70 |
| **MFQ** | -0.03 | 0.85 | -0.03 | 0.97 | -0.11 | 0.71 | -0.16 | 0.88 |
| **Baseline Age** | -0.01 | 0.82 | -0.01 | 0.99 | 0.10 | 0.71 | 0.15 | 0.88 |
| **Intracranial Volume (ICV)** | -0.81 | 1.18 | -0.68 | 0.50 | -1.07 | 1.05 | -1.02 | 0.31 |
| **Brain** | 1.58 | 114.78 | 0.01 | 0.99 | 2.18 | 118.72 | 0.02 | 0.99 |
| **Age-linear x Brain** | **-57.04** | **24.04** | **-2.37** | **0.02** | **-60.74** | **24.99** | **-2.43** | **0.02** |
| **Age-quadratic x Brain** | -11.91 | 22.47 | -0.53 | 0.60 | 11.99 | 21.43 | 0.56 | 0.58 |
| **Age-linear x Sex** | -16.07 | 23.59 | -0.68 | 0.50 | -19.74 | 20.14 | -0.98 | 0.33 |
| **Age-linear x W** | -6.29 | 44.08 | -0.14 | 0.89 | -5.61 | 35.53 | -0.16 | 0.87 |
| **Age-linear x H** | 2.52 | 42.77 | 0.06 | 0.95 | 3.35 | 34.36 | 0.10 | 0.92 |
| **Age-linear x High Risk** | -12.31 | 20.19 | -0.61 | 0.54 | -6.61 | 18.37 | -0.36 | 0.72 |
| **Age-linear x GenPop** | -24.67 | 44924.09 | 0.00 | 1.00 | -3.90 | 17124.75 | 0.00 | 1.00 |
| **Age-linear x SLES** | -9.99 | 24.93 | -0.40 | 0.69 | -15.30 | 21.04 | -0.73 | 0.47 |
| **Age-linear x SES** | 25.87 | 21.32 | 1.21 | 0.23 | 26.53 | 17.57 | 1.51 | 0.13 |
| **Age-linear x CTQ** | 1.92 | 19.66 | 0.10 | 0.92 | 4.47 | 17.83 | 0.25 | 0.80 |
| **Age-linear x Tanner** | 16.91 | 19.24 | 0.88 | 0.38 | 15.46 | 18.07 | 0.86 | 0.39 |
| **Age-linear x MFQ** | -11.20 | 19.34 | -0.58 | 0.56 | -12.20 | 16.73 | -0.73 | 0.47 |
| **Age-linear x Baseline Age** | 10.07 | 21.73 | 0.46 | 0.64 | 3.79 | 18.30 | 0.21 | 0.84 |
| **Age-linear x ICV** | 20.43 | 28.01 | 0.73 | 0.47 | 21.73 | 25.25 | 0.86 | 0.39 |
| **Age-quadratic x Sex** | 12.71 | 18.66 | 0.68 | 0.50 | 20.88 | 15.54 | 1.34 | 0.18 |
| **Age-quadratic x W** | 15.10 | 30.63 | 0.49 | 0.62 | 4.74 | 24.89 | 0.19 | 0.85 |
| **Age-quadratic x H** | 6.69 | 27.90 | 0.24 | 0.81 | 1.36 | 22.93 | 0.06 | 0.95 |
| **Age-quadratic x High Risk** | 18.42 | 15.16 | 1.21 | 0.22 | 12.60 | 13.16 | 0.96 | 0.34 |
| **Age-quadratic x GenPop** | 51.69 | 32354.53 | 0.00 | 1.00 | 41.45 | 13636.19 | 0.00 | 1.00 |
| **Age-quadratic x SLES** | -2.61 | 20.93 | -0.12 | 0.90 | 0.63 | 17.27 | 0.04 | 0.97 |
| **Age-quadratic x SES** | -7.67 | 16.53 | -0.46 | 0.64 | -10.84 | 13.83 | -0.78 | 0.43 |
| **Age-quadratic x CTQ** | 1.29 | 17.12 | 0.08 | 0.94 | -2.24 | 15.00 | -0.15 | 0.88 |
| **Age-quadratic x Tanner** | -2.06 | 15.21 | -0.14 | 0.89 | 0.47 | 13.83 | 0.03 | 0.97 |
| **Age-quadratic x MFQ** | -20.32 | 16.51 | -1.23 | 0.22 | -18.46 | 14.09 | -1.31 | 0.19 |
| **Age-quadratic x Baseline Age** | -11.03 | 15.32 | -0.72 | 0.47 | -5.90 | 13.37 | -0.44 | 0.66 |
| **Age-quadratic x ICV** | 18.31 | 21.99 | 0.83 | 0.41 | 7.89 | 20.98 | 0.38 | 0.71 |
| **Sex x Brain** | -0.42 | 0.76 | -0.55 | 0.58 | -0.76 | 0.67 | -1.14 | 0.25 |
| **W x Brain** | -0.07 | 1.10 | -0.06 | 0.95 | 0.39 | 0.89 | 0.43 | 0.67 |
| **H x Brain** | 0.25 | 1.06 | 0.23 | 0.81 | 0.52 | 0.87 | 0.59 | 0.56 |
| **High Risk x Brain** | -0.01 | 0.59 | -0.01 | 0.99 | 0.31 | 0.52 | 0.60 | 0.55 |
| **GenPop x Brain** | 1.95 | 893.93 | 0.00 | 1.00 | 2.23 | 924.63 | 0.00 | 1.00 |
| **SLES x Brain** | 0.34 | 0.60 | 0.57 | 0.57 | -0.08 | 0.53 | -0.16 | 0.88 |
| **SES x Brain** | 0.64 | 0.64 | 1.00 | 0.32 | 0.67 | 0.54 | 1.25 | 0.21 |
| **CTQ x Brain** | -0.12 | 0.66 | -0.18 | 0.86 | -0.11 | 0.57 | -0.20 | 0.84 |
| **Tanner x Brain** | 0.52 | 0.65 | 0.80 | 0.43 | 0.25 | 0.58 | 0.44 | 0.66 |
| **MFQ x Brain** | -0.20 | 0.70 | -0.28 | 0.78 | -0.15 | 0.58 | -0.26 | 0.79 |
| **Baseline Age x Brain** | 0.73 | 0.68 | 1.07 | 0.29 | 0.21 | 0.51 | 0.41 | 0.68 |
| **ICV x Brain** | -0.83 | 0.77 | -1.09 | 0.28 | -1.00 | 0.68 | -1.48 | 0.14 |

Volume of the right middle and superior frontal cortices are both associated with future initiation of alcohol consumption (p-fdr=0.035, 0,035). Standardized effects and uncorrected p-values are presented.

.

**Supplemental Table 5. Comparison of samples**

|  | **DNS (N=1303)** | **HCP (N=897)** | **TAOS^+ (^N=223)** | **t, F, *χ^2^*** | ***p*** |
| --- | --- | --- | --- | --- | --- |
| **Age** | 19.7 (1.25) 18-22 | 28.82 (3.68) 22-37 | 13.42 (0.96) 11.61-15.33 | F = 5420.39 | <1x10^-300^ |
| **Sex (% Female)** | 747 (57.33%) | 504 (56.19%) | 96 (43.05%) | *χ2* ***=*** 15.93 | **0.000347** |
| **m/AUDIT-C** | 3.76 (2.64) 0-11.69 | 3.42 (2.65) 0-11.38 | 1.26 (1.95) 0-9 | F = 89.43 | **3.38x10^-38^** |
| **European/ European American** | 579 (44.44%) | 597 (66.56%) | 126 (56.5%) | *χ2* ***=*** 105.31 | **1.35x10^-23^** |
| **African/African American** | 148 (11.36%) | 155 (17.28%) | 3 (1.35%) | *χ2* ***=*** 45.22 | **1.51x10^-10^** |
| **Asian/Asian American** | 353 (27.09%) | 45 (5.02%) | 4 (1.79%) | *χ2* ***=*** 225.94 | **8.68x10^-50^** |
| **Hispanic** | 83 (6.37%) | 73 (8.14%) | 73 (32.74%) | *χ2* ***=*** 157.54 | **6.18x10^-35^** |
| **Multi-racial/Native American/Other** | 140 (10.74%) | 27 (3.01%) | 17 (7.62%) | *χ2* ***=*** 45.29 | **1.46x10^-10^** |
| **CTQ** | 33.39 (8.12) 25-59.81 | - | 32.98 (7.09) 25-64 | t = 0.5 | 0.478993 |
| **PSS** | 14.65 (6.05) 0-32.89 | 13.07 (5.76) 0-35 | - | t = 37.59 | **1.03x10^-09^** |

Comparisons of measures that overlap across samples (DNS, HCP, TAOS). Continuous variables are presented in the format of: Mean (Standard deviation) Range. Dichotomous variables show the number of participants in that category, and the total percent of the sample that they comprise. Variables were winsorized prior to comparisons.

^+^= TAOS comparisons of mAUDIT-C scores were done with the maximum score attained by each participant.

-=Variable not present in sample.

DNS = Duke Neurogenetics Study; HCP = Human Connectome Project; TAOS = Teen Alcohol Outcomes Study; AUDIT = Alcohol Use Disorder Identification Test; PSS = Perceived Stress Scale; CTQ = Childhood Trauma Questionnaire.

**Supplemental Table 6.** **Frequency of Psychiatric Diagnosis in the DNS and HCP samples.**

| **Dataset** | **Disorder** | **N** |
| --- | --- | --- |
| DNS | Any | 258 |
|  | Depression | 66 |
|  | Bipolar | 35 |
|  | Anxiety | 50 |
|  | OCD | 16 |
|  | PTSD | 2 |
|  | Alcohol Abuse | 143 |
|  | Substance Abuse | 47 |
|  | Eating Disorder | 11 |
|  | Autism Spectrum | 1 |
|  | Psychosis | 3 |
| HCP | Any | 311 |
|  | Agoraphobia | 55 |
|  | Panic Disorder | 53 |
|  | Depression | 75 |
|  | Alcohol Abuse/Dependence | 176 |
|  | Marijuana Abuse/Dependence | 103 |

80 participants in the DNS have more than one diagnosis.

103 participants in the HCP have more than one diagnosis.

**Supplemental Table 7. Comparison of DNS subjects who did or did-not complete follow-up questionnaires**

|  | **No Follow-Up (N=624)** | **Follow-Up (N=679)** | **t/χ^2^** | ***p*** |
| --- | --- | --- | --- | --- |
| **AUDIT-C** | 4.0492 (2.6632) | 3.4993 (2.595) | 3.7694 | **0.0002** |
| **AUDIT Total** | 5.5913 (4.4163) | 4.8778 (4.1872) | 2.9869 | **0.0029** |
| **Age** | 19.8125 (1.2739) | 19.592 (1.2265) | 3.1765 | **0.0015** |
| **P-SES** | 7.5065 (1.6722) | 7.3884 (1.7094) | 1.2603 | 0.2078 |
| **P-ED** | 7.0232 (1.6025) | 7.0751 (1.7291) | -0.562 | 0.5742 |
| **PSS** | 14.4852 (5.9435) | 14.7976 (6.1404) | -0.9328 | 0.3511 |
| **CTQ** | 33.7902 (8.3233) | 33.016 (7.9077) | 1.7179 | 0.0861 |
| **Frontal Volume** | 0.4834 (0.0559) | 0.4815 (0.0564) | 0.6047 | 0.5455 |
| **Insula Volume** | 0.5498 (0.0539) | 0.5505 (0.0547) | -0.2484 | 0.8038 |
| **ICV** | 1.4877 (0.1451) | 1.4754 (0.1384) | 1.5573 | 0.1196 |
| **Sex (number of female respondents)*** | N=320 (51.28%) | N=427 (62.89%) | 17.429 | **2.98x10^-05^** |
| **Diagnosis (non substance-related) *** | N=77 (12.34%) | N=72 (10.6%) | 0.8037 | 0.3700 |
| **Caucasian*** | N=253 (40.54%) | N=326 (48.01%) | 7.0435 | **0.0080** |
| **African/African American*** | N=86 (13.78%) | N=62 (9.13%) | 6.5319 | **0.0106** |
| **Asian/Asian American*** | N=171 (27.4%) | N=182 (26.8%) | 0.0327 | 0.8564 |
| **Hispanic*** | N=42 (6.73%) | N=41 (6.04%) | 0.1582 | 0.6908 |
| **Multi-racial/Native American/Other*** | N=72 (11.54%) | N=68 (10.01%) | 0.6364 | 0.4250 |

*=Analysis was run as a chi-squared test, all others were run as t-tests.

AUDIT = Alcohol Use Disorder Identification Test; CTQ = Childhood Trauma Questionnaire; P-ED = Parental Education; P-SES = Perceived Socioeconomic Status; PSS = Perceived Stress Scale. ; ICV = Intracranial Volume

**Supplemental Table 8. Association of Alcohol Consumption/Initiation with Covariates**

| **Study** | **DNS** |  | **HCP** |  | **TAOS*** |  |
| --- | --- | --- | --- | --- | --- | --- |
| **Variable** | **statistic (t, F, r)** | ***p*** | **statistic (t, F, r)** | ***p*** | **statistic (t, *χ^2^* )** | ***p*** |
| Sex (Female vs Male) | t = 9.1624 | **2.70x10^-19^** | t = 8.7447 | **1.72x10^-17^** | *χ2* ***=*** 6.24x10^-31^ | 1.0000 |
| Diagnosis (non Substance-related) (vs none) | t = -0.3185 | 0.7505 | t = -0.6746 | 0.5007 | **-** | **-** |
| Ethnicity | F = 29.7684 | **5.82x10^-8^** | F = 2.6516 | 0.1038 | *χ2* **=** 6.5217 | 0.5890 |
| European/ European American (vs not) | t = -7.6998 | **2.84x10^-14^** | t = -3.2474 | **0.0012** | *χ2* ***=*** 0.0424 | 0.8368 |
| African/African American (vs not) | t = 3.1538 | **0.0019** | t = 4.1931 | **3.89x10^-5^** | **-** | **-** |
| Asian/Asian American (vs not) | t = 5.0911 | **4.54x10^-7^** | t = 1.4615 | 0.1502 | **-** | **-** |
| Hispanic (vs not) | t = -0.0295 | 0.9765 | t = -0.9598 | 0.3398 | *χ2* ***=*** 0.0908 | 0.7631 |
| Multi-racial/Native American/Other (vs not) | t = 2.2086 | **0.0285** | t = -0.3249 | 0.7477 | **-** | **-** |
| Age | r = 0.126 | **5.10x10^-6^** | r = -0.1222 | **0.0002** | t = -2.8750 | **0.0045** |
| P-SES/SES | r = 0.1747 | **2.15x10^-10^** | r = 0.0102 | 0.7604 | t = -1.7684 | 0.0789 |
| PSS/SLES | r = -0.0383 | 0.1669 | r = 0.023 | 0.4908 | t = -0.8601 | 0.3908 |
| P-ED/ED | r = 0.0883 | **0.0014** | r = -0.0153 | 0.6465 | **-** | **-** |
| CTQ | r = -0.0966 | **0.0005** | **-** | **-** | t = -1.6842 | 0.0945 |
| Scanner (1 vs 2) | t = -0.5507 | 0.5822 | **-** | **-** | **-** | **-** |
| High-risk for depression (TAOS-only) | **-** | **-** | **-** | **-** | t = 0.5780 | 0.4471 |
| Tanner stage | **-** | **-** | **-** | **-** | t = -2.0118 | **0.0459** |
| MFQ | **-** | **-** | **-** | **-** | t = 0.2417 | 0.8094 |

Association of model covariates with alcohol consumption (AUCIT-C/mAUDIT-C).

* = For analyses in TAOS, comparisons are between baseline measurements of participants who initiate, and those who do not.

Comparisons were run as t-tests, anovas, pearson correlations, or chi-squared tests. SES = Socioeconomic status, PSS = Perceived Stress Scale, ED = education, CTQ = Childhood Trauma Questionnaire, MFQ = Mood and Feelings Questionnaire; SLES = Stressful Life Events Schedule. DNS = Duke Neurogenetics Study; HCP = Human Connectome Project; TAOS = Teen Alcohol Outcomes Study

- = Not present in sample, not applicable, or insufficient numbers to run test.

**Supplemental Table 9: Brain volume does not predict impulsivity, negative urgency, or intelligence.**

|  |  | **DNS** |  |  |  |  | **HCP** |  |  |  |  |
| --- | --- | --- | --- | --- | --- | --- | --- | --- | --- | --- | --- |
| **Y** | **X** | **Beta** | **Δ r^2^** | **P** | **P-fdr** | **N** | **Beta** | **Δ r^2^** | **P** | **P-fdr** | **N** |
| Coping | Frontal | -0.0876 | 0.0033 | 0.0293 | 0.1529 | 1303 | - | - | - | - | - |
|  | Insula | -0.0915 | 0.0036 | 0.0237 | 0.1529 | 1303 | - | - | - | - | - |
| Self-report Impulsivity | Frontal | 0.0487 | 0.0010 | 0.1909 | 0.2864 | 1303 | -0.0844 | 0.0053 | 0.0103 | 0.0824 | 897 |
|  | Insula | 0.0680 | 0.0020 | 0.0692 | 0.1529 | 1303 | -0.0549 | 0.0046 | 0.1045 | 0.3739 | 897 |
| IQ | Frontal | 0.0760 | 0.0025 | 0.0514 | 0.1529 | 1275 | 0.0060 | -5.24E-05 | 0.8295 | 0.9271 | 897 |
|  | Insula | 0.0699 | 0.0021 | 0.0764 | 0.1529 | 1275 | -0.0026 | -2.71E-05 | 0.9271 | 0.9271 | 897 |
| DDT - K | Frontal | -0.0891 | 0.0034 | 0.0436 | 0.1529 | 1157 | -0.0142 | 3.62E-05 | 0.6877 | 0.9169 | 891 |
|  | Insula | -0.0320 | 0.0004 | 0.4720 | 0.5664 | 1157 | 0.0534 | -0.0089 | 0.1402 | 0.3739 | 891 |
| NEO - N5 | Frontal | -0.0636 | 0.0018 | 0.1024 | 0.1756 | 1302 | - | - | - | - | - |
|  | Insula | -0.0478 | 0.0010 | 0.2232 | 0.2976 | 1302 | - | - | - | - | - |
| NEO - N | Frontal | -0.0008 | 2.64E-07 | 0.9789 | 0.9789 | 1302 | -0.0212 | -0.0001 | 0.3848 | 0.6157 | 891 |
|  | Insula | -0.0080 | 2.73E-05 | 0.7879 | 0.8595 | 1302 | -0.0308 | 0.0014 | 0.2337 | 0.4674 | 891 |

Parameters from linear regression models of brain volume predicting behavior. Covariates were the same as in whole-brain analyses. Coping = substance-use subscale of the brief COPE. Self-report impulsivity: DNS - Barratt Impulsiveness Scale; HCP - Achenbach Adult Self-Report Impulsivity. IQ: DNS: WASI-II; HCP: NIH Toolbox. DDT – K: Delayed discounting task. NEO-N: Neuroticism. NEO-N5: Neuroticism Impulsivity subscale. Standardized effect-sizes (ie beta values) are presented. Δ r^2^: Change in model-fit when the x-variable is added (negative values indicate the model-fit decreased).

**Supplemental Table 10: Brain volume clusters do not predict impulsivity, negative urgency, or intelligence.**

| **Study** | **Y** | **X** | **Beta** | **Δ r^2^** | **P** | **P-fdr** | **N** |
| --- | --- | --- | --- | --- | --- | --- | --- |
| **DNS** | BCOPE - Sub | Insula_48_6_5 | -0.0767 | 0.0044 | 0.0127 | 0.1110 | 1303 |
|  |  | MidFrontal_27_39_26 | -0.0888 | 0.0048 | 0.0089 | 0.1110 | 1303 |
|  |  | MidFrontal_38_18_51 | -0.0291 | 0.0006 | 0.3762 | 0.5424 | 1303 |
|  |  | SupFrontal_29_62_8 | -0.0706 | 0.0030 | 0.0392 | 0.1880 | 1303 |
|  | BIS | Insula_48_6_5 | 0.0313 | 0.0007 | 0.2726 | 0.5424 | 1303 |
|  |  | MidFrontal_27_39_26 | 0.0402 | 0.0010 | 0.2010 | 0.4825 | 1303 |
|  |  | MidFrontal_38_18_51 | -0.0006 | 2.44x10^-07^ | 0.9840 | 0.9924 | 1303 |
|  |  | SupFrontal_29_62_8 | 0.0325 | 0.0006 | 0.3053 | 0.5424 | 1303 |
|  | IQ | Insula_48_6_5 | 0.0232 | 0.0004 | 0.4388 | 0.5850 | 1275 |
|  |  | MidFrontal_27_39_26 | 0.0793 | 0.0038 | 0.0163 | 0.1110 | 1275 |
|  |  | MidFrontal_38_18_51 | 0.0137 | 0.0001 | 0.6669 | 0.8003 | 1275 |
|  |  | SupFrontal_29_62_8 | 0.0329 | 0.0007 | 0.3219 | 0.5424 | 1275 |
|  | DDT - k | Insula_48_6_5 | -0.0315 | 0.0007 | 0.3509 | 0.5424 | 1157 |
|  |  | MidFrontal_27_39_26 | -0.0223 | 0.0003 | 0.5521 | 0.6974 | 1157 |
|  |  | MidFrontal_38_18_51 | -0.0321 | 0.0007 | 0.3709 | 0.5424 | 1157 |
|  |  | SupFrontal_29_62_8 | -0.0644 | 0.0025 | 0.0864 | 0.2593 | 1157 |
|  | NEO - N5 | Insula_48_6_5 | -0.0441 | 0.0014 | 0.1396 | 0.3721 | 1302 |
|  |  | MidFrontal_27_39_26 | -0.0287 | 0.0005 | 0.3842 | 0.5424 | 1302 |
|  |  | MidFrontal_38_18_51 | -0.0749 | 0.0037 | 0.0185 | 0.1110 | 1302 |
|  |  | SupFrontal_29_62_8 | -0.0646 | 0.0025 | 0.0515 | 0.2059 | 1302 |
|  | NEO - N | Insula_48_6_5 | 0.0063 | 2.98x10^-05^ | 0.7788 | 0.8608 | 1302 |
|  |  | MidFrontal_27_39_26 | 0.0002 | 3.45x10^-08^ | 0.9924 | 0.9924 | 1302 |
|  |  | MidFrontal_38_18_51 | -0.0431 | 0.0012 | 0.0730 | 0.2501 | 1302 |
|  |  | SupFrontal_29_62_8 | 0.0067 | 2.71x10^-05^ | 0.7890 | 0.8000 | 1302 |
| **HCP** | DDT - k | Frontal_20_24_42 | 0.0545 | -0.0058 | 0.1084 | 0.4279 | 891 |
|  |  | Insula_38_12_0 | 0.0258 | -0.0012 | 0.4520 | 0.5668 | 891 |
|  | ASR_IMP | Frontal_20_24_42 | -0.0453 | 0.0008 | 0.1605 | 0.4279 | 897 |
|  |  | Insula_38_12_0 | -0.0328 | 0.0034 | 0.3124 | 0.5668 | 897 |
|  | NEO - N | Frontal_20_24_42 | 0.0166 | -0.0003 | 0.4960 | 0.5668 | 891 |
|  |  | Insula_38_12_0 | -0.0198 | 0.0010 | 0.4195 | 0.5668 | 891 |
|  | IQ | Frontal_20_24_42 | -0.0446 | -0.0005 | 0.0747 | 0.4279 | 897 |
|  |  | Insula_38_12_0 | -0.0115 | 1.84x10^-05^ | 0.6506 | 0.6506 | 897 |

Parameters from linear regression models of brain volume predicting behavior. Covariates were the same as in whole-brain analyses. BCOPE - Sub = substance-use subscale of the brief COPE. BIS = Barratt Impulsiveness Scale; ASR IMP = Achenbach Adult Self-Report Impulsivity. IQ: DNS: WASI-II; HCP: NIH Toolbox. DDT – K: Delayed discounting task. NEO-N: Neuroticism. NEO-N5: Neuroticism Impulsivity subscale. Standardized effect-sizes (ie beta values) are presented. Δ r^2^: Change in model-fit when the x-variable is added (negative values indicate the model-fit decreased).
